# Binary decision between asymmetric and symmetric cell division is defined by the balance of PAR proteins in *C. elegans* embryos

**DOI:** 10.1101/2020.08.24.264135

**Authors:** Yen Wei Lim, Fu-Lai Wen, Prabhat Shankar, Tatsuo Shibata, Fumio Motegi

**Author notes:** These authors contributed equally (Fu-Lai Wen, Prabhat Shankar). Correspondence should be addressed to T.S. or F.M.

## Abstract

Coordination between cell differentiation and proliferation during development requires the balance between asymmetric and symmetric modes of cell division. However, the cellular intrinsic cue underlying the binary choice between these two division modes remains elusive. Here we show evidence in *Caenorhabditis elegans* that the invariable lineage of the division modes is programmed by the balance between antagonizing complexes of partitioning-defective (PAR) proteins. By uncoupling unequal inheritance of PAR proteins from that of fate determinants during zygote division, we demonstrated that changes in the balance between PAR-2 and PAR-6 are sufficient to re-program the division modes from symmetric to asymmetric and *vice versa* in two-cell stage embryos. The division mode adopted occurs independently of asymmetry in cytoplasmic fate determinants, cell-size asymmetry, and cell-cycle asynchrony between the sister cells. We propose that the balance between antagonizing PAR proteins represents an intrinsic self-organizing cue for binary specification of the division modes during development.

## INTRODUCTION

The development of a multicellular organism requires strict control of the balance between cell proliferation and differentiation. Within a developing embryo, each cell proliferates via either asymmetric or symmetric mode of cell division (Horvitz and Herskowitz, 1992), yielding two daughter cells with either distinct or identical cellular constituents, respectively. The symmetric mode of cell division gives rise to two copies of the mother cell. The asymmetric mode of cell division results in unequal inheritance of fate determinants, which will promote the subsequent induction of distinct cellular fates between the daughter cells. A feature of asymmetric cell division can be accompanied by differences in cellular size between the daughter cells, although the cellular size asymmetry is not necessary for induction of distinct cellular fates during asymmetric cell division. The balance between asymmetric and symmetric modes of cell division is controlled during embryogenesis as well as during postnatal development. Many adult stem cells exhibit the homeostatic control of self-renewal and differentiation by balancing the two modes of cell division (Chen et al., 2016; Knoblich, 2008). Disruption of this balance results in developmental disorders, premature depletion of the stem cells, and abnormal growth of prenatal and postnatal organs.

The binary specification between asymmetric and symmetric modes of cell division is regulated by an intrinsic program (i.e., cell-autonomous manner) and by extrinsic stimuli that influence the intrinsic program (i.e., non-cell-autonomous manner). The resultant decision between the two division modes is thought to control the distribution of cell-polarity regulators, which in turn directs the spatial organization of the cell (Morrison and Kimble, 2006). Therefore, these cell-polarity regulators have been considered as passive effectors of the intrinsic cell-division program. To date, many machineries have been deemed essential for either symmetric or asymmetric mode of cell division. However, it has been challenging to obtain rigorous data identifying the nature of the intrinsic program used to impose the binary choice between the two division modes. Indeed, many arguments in support of an apparent intrinsic program have relied on circumstantial evidence, such as a combination of the mother cell polarity, spatial asymmetry in the external environment, and unequally sized daughter cells. Thus, the identity of the cue that dictates to the intrinsic cell-division program is not completely understood.

We investigated the intrinsic program underlying the binary specification between the two modes of cell division using the classic example of invariant cell-division lineage in *C. elegans* (Sulston and Horvitz, 1977; Sulston et al., 1983). The newly-fertilized *C. elegans* zygote undergoes asymmetric cell division in a cell-autonomous manner, yielding two daughter cells, AB and P1 (Figure 1). During the first asymmetric cell division, the inner layer of the plasma membrane (i.e. the cell cortex) is compartmentalized via segregation of the conserved cell-polarity regulators, PAR proteins (Goldstein and Macara, 2007; Kemphues, 2000). PAR-3, PAR-6, atypical protein kinase C (aPKC), and the active form of Cdc42 GTPase become enriched at the anterior cortex (Etemad-Moghadam et al., 1995; Hung and Kemphues, 1999; Tabuse et al., 1998), whereas PAR-1 and PAR-2 localize at the posterior cortex (Boyd et al., 1996; Guo and Kemphues, 1995) (Figure 1). It has been widely accepted that cortical patterning of PAR proteins is largely attributed to the principle of mutual inhibition, wherein proteins from one cortical domain antagonize the co-localization of proteins from the other domain and *vice versa* (Hoege and Hyman, 2013). These spatially-biased PAR proteins in turn segregate cytoplasmic fate determinants (e.g., MEX-5, PIE-1, and P-granules) and displace the mitotic spindle toward the posterior pole. Consequently, the two daughter cells comprise distinct concentration of fate determinants, cell size asymmetry and cell-cycle asynchrony (Griffin, 2015; Rose and Gonczy, 2014) (Figure 1). Displacement of the mitotic spindle and the consequent asymmetry in cellular size are not required to ensure the unequal inheritance of the fate determinants (Gotta and Ahringer, 2001; Gotta et al., 2003). In this study, asymmetric and symmetric modes of cell division were defined as the cellular ability to accomplish asymmetric segregation and unequal inheritance of both PAR proteins and fate determinants during mitosis.

**Figure 1.**
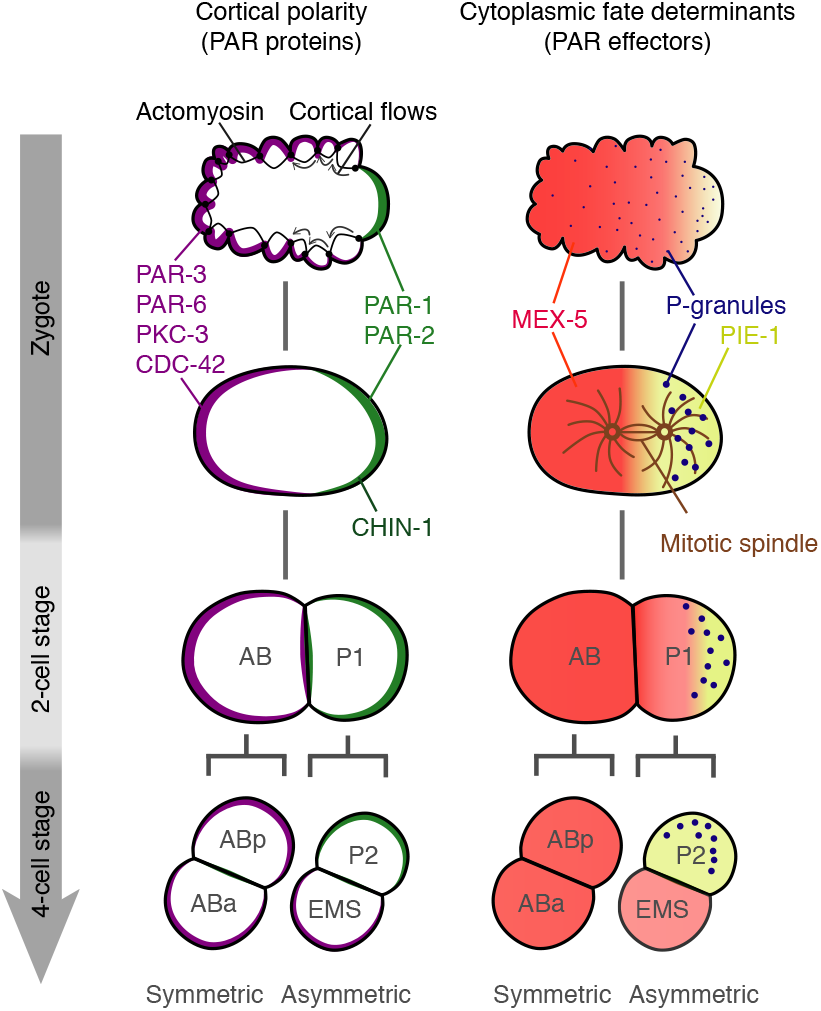
A schematic of *C. elegans* embryo polarization. Prior to polarization, a zygote localizes PAR-3, PAR-6, PKC-3, and CDC-42 throughout the cortex, and distributes PAR-1, PAR-2, and CHIN-1 in the cytoplasm. During symmetry breaking, the advective flows of cortical actomyosin networks translocate PAR-3, PAR-6, PKC-3, and CDC-42 to the anterior cortical domain. Active PKC-3 kinase at the cortex excludes PAR-1, PAR-2, and CHIN-1 from the anterior cortical domain, allowing them to localize onto the posterior cortical domain. Asymmetries in these PAR proteins in turn mediate the segregation of their effectors, including P-granules, MEX-5, PIE-1, and the mitotic spindle, during mitosis. In two-cell stage embryos, the anterior AB cell is destined to divide symmetrically, resulting in equal inheritance of PAR proteins and fate determinants by the two daughter cells, ABa and ABp. In contrast, the posterior P_1_ cell is engaged to undergo asymmetric cell division, resulting in unequal inheritance of PAR proteins and fate determinants by the two daughter cells, EMS and P2. All zygotes and embryos are oriented with the posterior to the right in this and all subsequent figures.

The above-described cell polarity cascade in zygotes dictates the two daughter cells, AB and P1, to undergo symmetric and asymmetric cell division, respectively. During the second cell division, the polarized distributions of cortical PAR proteins and the cytoplasmic fate determinants are reiterated only in P1 cells, but not in AB cells (Figure 1). This commitment of AB and P1 cells to their respective mode of cell division occurs in a cell-autonomous manner (Priess and Thomson, 1987), indicating that the cue that dictates to the intrinsic cell-division program must be unequally inherited during the first cell division. AB and P1 cells also acquire differential abilities regarding the rotation of mitotic spindles. The AB spindle remains orthogonal to the anteroposterior axis, whereas the P1 spindle is rotated and aligned with the axis. In contrast to the cell-autonomous specification of the division modes, this rotation of mitotic spindles is directed by a combination of cell-autonomous manner [PAR proteins (Cheng et al., 1995; Kemphues et al., 1988) and microtubule-pulling forces (Kotak, 2019)] and non-cell-autonomous manner [the mid-body structure (Singh and Pohl, 2014) and a physical contact between AB and P1 cells (Sugioka and Bowerman, 2018)]. Still, the identity of the cue dictating the decision between symmetric and asymmetric modes of cell division in AB and P1 cells remains unclear.

In this report, we developed experimental systems that enabled independent assessment of the roles of PAR proteins and those of asymmetries in fate determinants, cell size and cell-cycle progression in two-cell stage embryos. Using genetic manipulations that uncoupled unequal inheritance of PAR proteins from that of the fate determinants during the first cell division, we demonstrated that changes in the balance between antagonizing PAR proteins are sufficient to re-program the cell division mode from symmetric to asymmetric and *vice versa* in two-cell stage embryos. Moreover, the specification of a cell division mode occurs independently of the unequal inheritance of fate determinants, cell size asymmetry, and cell-cycle asynchrony between the sister cells. Therefore, we propose that the balance between PAR proteins represents an intrinsic self-organizing cue for binary specification regarding the cell-division modes.

## RESULTS

### Manipulation of PAR protein inheritance during the first cell division

To isolate the roles of PAR proteins from those of other asymmetric features in the specification of the two division modes, we sought to control the amount of PAR proteins in AB and P1 cells without affecting asymmetries in the fate determinants, cellular size, and cell-cycle progression in two-cell stage embryos. From the one-cell to the two-cell stage, transcriptional and translational regulation generally remains quiescent (Seydoux and Dunn, 1997; Seydoux et al., 1996). Accordingly, conventional transgene-based approaches cannot be used for the efficient manipulation of the levels and activities of PAR proteins from the onset of the second cell division. Therefore, we attempted to modify the PAR protein levels in AB and P1 cells by manipulating inheritance of PAR proteins during the first cell division. The prevailing model of PAR protein patterning involves reciprocal exclusion at the cortex, wherein the cortical concentration of each protein is determined by the spatially-biased rates of cortical association and disassociation in the respective domains (Blanchoud et al., 2015; Cuenca et al., 2003; Dawes and Munro, 2011; Goehring et al., 2011; Hoege and Hyman, 2013). The expansion of one cortical PAR domain is antagonized not only by the opposing PAR proteins but also by a limiting pool of maternally-supplied PAR proteins (Goehring et al., 2011). We thus attempted to modify the levels of specific PAR proteins in zygotes by altering the corresponding maternal supplies.

We first manipulated the maternal supplies of PAR-2 and PAR-6 via transgene expressions in germline. The germline expression of GFP::PAR-2 rescued the polarized distribution of mCherry::PAR-6 in *par-2(ok1723)* zygotes from which endogenous *par-2* had been deleted (Figures 2A and 2B). The germline expression of mCherry::PAR-6 restored the polarized localization of GFP::PAR-2 in *par-6(3’ UTR RNAi)* zygotes (Figure S1A). The germline levels of GFP::PAR-2 were modified by adapting the codon usage in the *gfp::par-2* transgene (Goehring et al., 2011; Redemann et al., 2014). A stepwise increase in the transgene codon-adaptation-index (CAI) from 0.26 to 0.60 resulted in a progressive increase in the cytoplasmic intensities of GFP::PAR-2 in zygotes (Figure S1B). Consistent with a previous report (Goehring et al., 2011), zygotes overexpressing *gfp::par-2* transgenes exhibited enlarged posterior PAR-2 domains compared with the control zygotes (Figures 2A and 2C). These enlarged posterior PAR-2 domains were restored to a normal size by the co-expression of mCherry::PAR-6 (Figures 2A and 2C). There results indicate that changes in the balance between PAR-2 and PAR-6 are sufficient to alter the size of the cortical PAR domains in zygotes. We then established a set of stable transgenic animals expressing various combinations of *gfp::par-2* transgenes (in the *par-2(ok1723)* background) and *mCherry::par-6* transgene (in the presence of endogenous *par-6*). These transgenes were combined with additional genetic mutation, either *lgl-1(dd21)* or *nos-3(q650)*, which respectively increases or reduces the level of maternally-supplied endogenous PAR-6 (Beatty et al., 2010; Hoege et al., 2010; Pacquelet et al., 2008). We then measured the sizes of the anterior cortical domain (mCherry::PAR-6) and the posterior cortical domain (GFP::PAR-2) in these zygotes at the steady state (i.e., during the maintenance phase).

**Figure 2.**
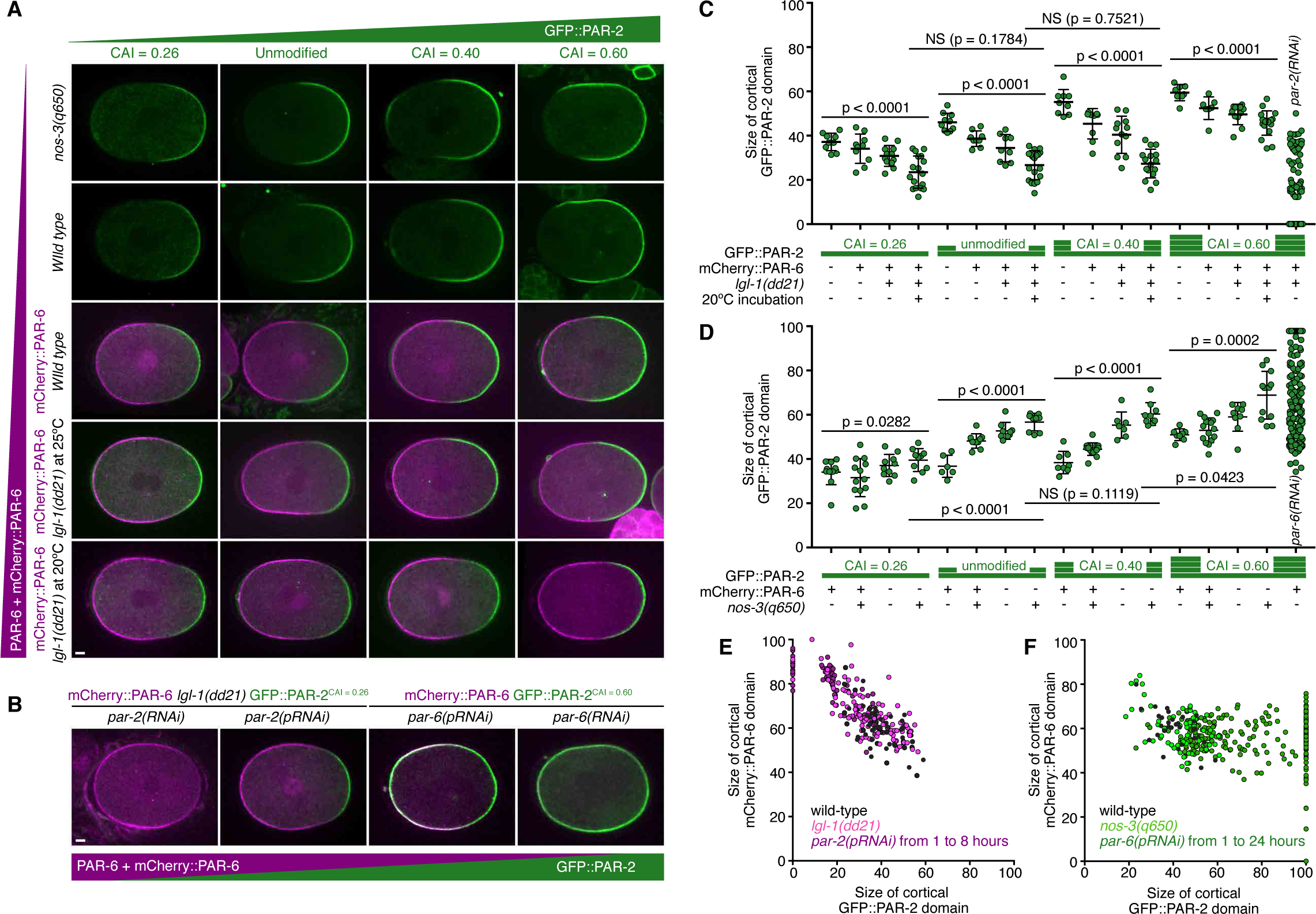
The landscape of PAR polarity patterning in *C. elegans* zygotes. (A)A balance between PAR-2 and PAR-6 levels defines the cortical pattern of PAR proteins in zygotes. PAR-2 levels were modified by means of *gfp::par-2* transgene codon adaptation (codon-adaptation index (CAI) values from 0.26 to 0.60) and of temperature (either at 25ºC or at 15ºC). PAR-6 levels were changed by means of *mCherry::par-6* transgene and of *nos-3(q650)* or *lgl-1(dd21)* mutation. Representative zygotes during the maintenance phase are shown. Scale Bar, 5 μm. (B) Defining the limit of PAR protein segregation. Representative images of *par-2(RNAi)* zygotes, *par-2(partial RNAi: pRNAi)* zygotes wherein PAR-6 was predominant, *par-6(RNAi)* zygotes, and *par-6(pRNAi)* zygotes wherein PAR-2 was predominant are shown. Scale Bar, 5 μm. (C and D) The graphs depict the size of GFP::PAR-2 cortical domains in zygotes with various PAR balances. Height of green bars indicate the *gfp::par-2* transgene CAI. (C) Data represent mean ± s.d. from n = 9, 10, 12, 17, 10, 9, 9, 19, 9, 9, 12, 16, 8, 7, 11, 19, 98 zygotes. (D) Data represent mean ± s.d. from n = 9, 11, 8, 8, 4, 8, 7, 7, 6, 13, 5, 7, 6, 13, 7, 10, 194 zygotes. *p*-values; Mann–Whitney test. (E and F) The sizes of mCherry::PAR-6 and GFP::PAR-2 cortical domains in zygotes with various balances of PAR proteins. Black circles indicate zygotes on wild-type background. Light purple (E) and light green (F) indicate zygotes on *lgl-1(dd21)* and *nos-3(q650)* backgrounds, respectively. Dark purple (E) and dark green (F) denote zygotes subjected to *par-2(pRNAi)* and *par-6(pRNAi)* treatments, respectively.

We first examined the patterns of cortical PAR domains in zygotes with a predominance of PAR-6. A stepwise reduction in the level of GFP::PAR-2 led to a progressive decrease in the size of the posterior PAR-2 domain (Figures 2A and 2C) and in a reciprocal increase in the size of the anterior PAR-6 domain (Figure S1C). The various sizes of the cortical PAR domains were converted to a consistent size (82% and 18% of the mCherry::PAR-6 and GFP::PAR-2 domains, respectively) by applying the same genetic treatment that enabled the predominance of PAR-6 (i.e., simultaneous increase in PAR-6 and decrease in GFP::PAR-2 levels) (Figures 2A and 2C). Any further reduction in the GFP::PAR-2 levels via partial *par-2(RNAi)* treatment [*par-2(pRNAi)*] led to the disappearance of the GFP::PAR-2 domain from the posterior cortex (Figures 2B, 2C, 2E, S1E, and S1G). We confirmed that the *par-2(pRNAi)* treatment did not completely eliminate GFP::PAR-2 from these zygotes, as indicated by the GFP::PAR-2 accumulation at the cell-cell contact in two-cell stage embryos. These observations suggest that the minimum size of the cortical PAR-2 domain is maintained by a threshold. Especially, the establishment of the posterior PAR domain exhibits a bi-stable switch-like characteristic, wherein the polarized state (i.e., the co-existent polarized PAR-6 and PAR-2 domains) can abruptly shift to an unpolarized state (i.e., homogeneous PAR-6 domain and no PAR-2 domain) if the posterior PAR domain size falls below the threshold.

We next applied a stepwise increase in the GFP::PAR-2 levels and reduction in the PAR-6 levels to yield zygotes in which PAR-2 is predominant. Remarkably, simultaneous increase in GFP::PAR-2 and decrease in PAR-6 levels caused the distribution of GFP::PAR-2 to extend into the anterior cortex, where mCherry::PAR-6 remained significantly enriched (Figures 2A, 2B, 2D, 2F, S1D, S1F, and S1H). A live-imaging analysis revealed that the boundary of the cortical mCherry::PAR-6 domain co-migrated with the posterior-to-anterior movement of cortical ruffling, whereas GFP::PAR-2 was seen throughout the cortex (Figures S2C, S2F, and S3). These results indicate that the PAR system appears to permit the co-existence of an anteriorly-polarized PAR-6 domain with a PAR-2 domain throughout the cortex. This violation of the reciprocal exclusion between the anterior and the posterior PAR domains cannot be explained by the conventional models of cortical PAR patterning, wherein the anterior group of proteins (PAR-3, PAR-6, PKC-3, and Cdc42) and the posterior group of proteins (PAR-1 and PAR-2) are segregated throughout the polarization processes (Blanchoud et al., 2015; Dawes and Munro, 2011; Goehring et al., 2011).

### Maintenance of polarized PAR domains via a combinatorial network of two reciprocal exclusion pathways

To understand how PAR-6 remains polarized in the absence of reciprocal exclusion with PAR-2, we investigated the distributions of other PAR proteins. In zygotes with a PAR-2 predominance, the polarized mCherry::PAR-6 domain was maintained, while GFP::PAR-2 and endogenous PAR-1 were localized throughout the cortex (Figures 3A-3D and S3A). However, endogenous PAR-3 was significantly minimized or was undetectable at the anterior cortex (Figures 3A, 3E, and S3B), suggesting that the polarized state of PAR-6 is maintained independently of cortical PAR-3 loading. PAR-3 is considered as a cortical scaffold that locally recruits PAR-6 and PKC-3 to the active form of CDC-42 at the anterior cortex (Rodriguez et al., 2017; Wang et al., 2017). Despite the essential functions of PAR-3 in wild-type zygotes, PAR-3 becomes dispensable for the cortical loading of PAR-6 and PKC-3 in zygotes depleted of the Hsp90 co-chaperone CDC-37 (Beers and Kemphues, 2006), suggesting another scaffold for PAR-6 and PKC-3. Because PAR-6 contains a CRIB domain that directly associates with the active form of CDC-42 (Aceto et al., 2006; Joberty et al., 2000), we hypothesized that active CDC-42 recruits PAR-6 directly to the anterior cortex in a manner independent on both cortical PAR-3 loading and antagonism between PAR-2 and PAR-6. To test this hypothesis, we altered the asymmetric distribution of active CDC-42 by depleting the CDC-42 GTPase-activating protein, CHIN-1. CHIN-1 localizes at the posterior cortex independently of other posterior PAR proteins, and restricts CDC-42 activation only at the anterior cortex (Kumfer et al., 2010; Sailer et al., 2015). In this study, the depletion of CHIN-1 from PAR-2-predominant zygotes did not affect the initial segregation of mCherry::PAR-6 toward the anterior cortical domain. However, these zygotes failed to maintain the anteriorly-enriched mCherry::PAR-6 domain after the cessation of cortical flows (Figures 3F, 3G, and S3C). Therefore, we conclude that the polarized state of PAR-6 is maintained by the accumulation of active CDC-42 at the anterior cortex through a process mediated by CHIN-1.

**Figure 3.**
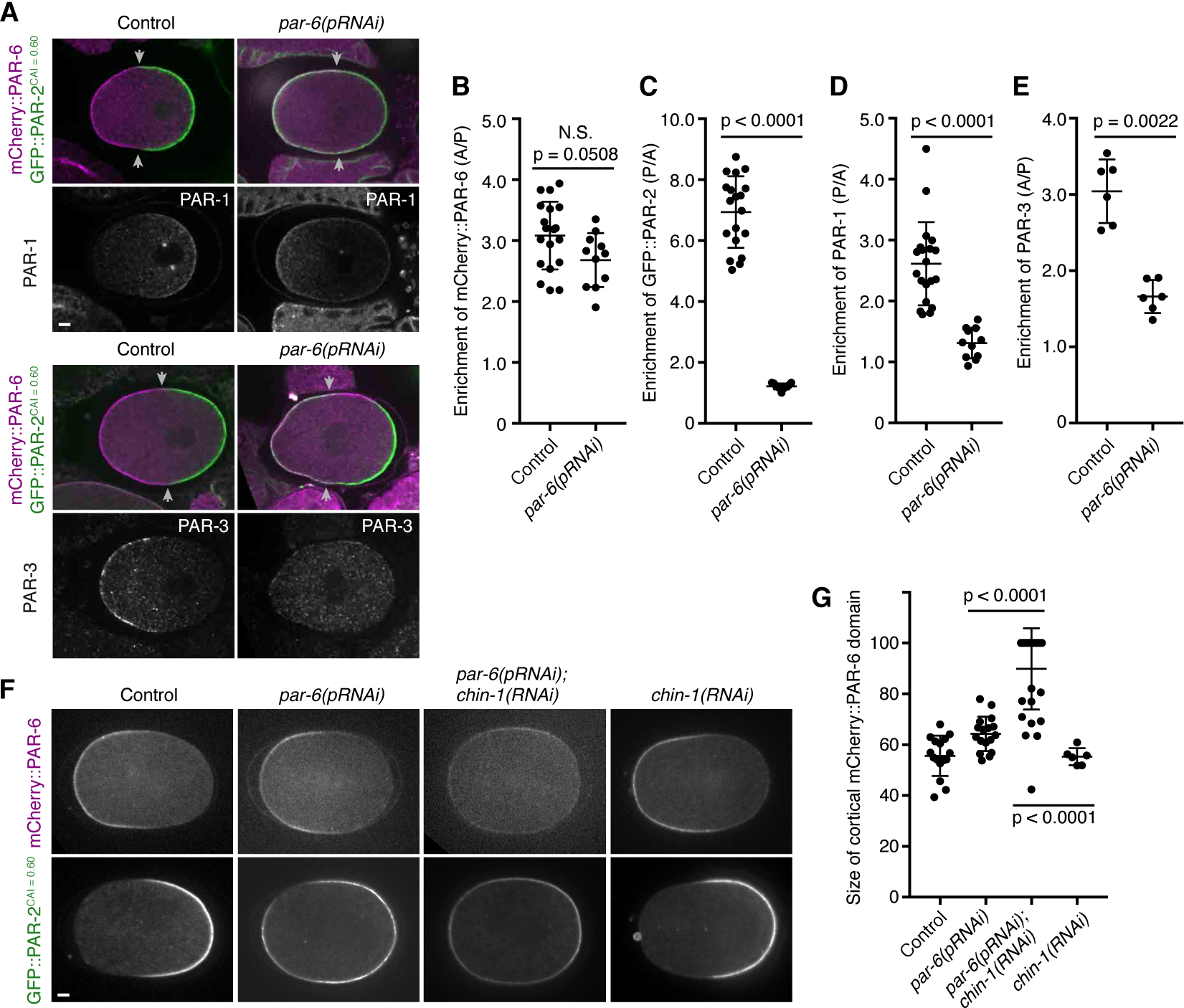
A combinatorial network of two reciprocal exclusion pathways ensures cortical polarization. (A) The polarized distribution of PAR-3 is not required for the maintenance of polarized PAR-6 domain. Representative images of mCherry::PAR-6, GFP::PAR-2, PAR-1, and PAR-3 in control zygotes and *par-6(pRNAi)* zygotes wherein PAR-2 was predominant are shown. Arrowheads show the edges of mCherry::PAR-6 cortical domain. Scale Bar, 5 μm. (B-E) The graphs depict the ratios of the cortical distribution of mCherry::PAR-6 (B), GFP::PAR-2 (C), PAR-1 (D), and PAR-3 (E) between the anterior and the posterior cortical domains in control and *par-6(pRNAi)* zygotes wherein PAR-2 was predominant. Data present mean ± s.d. from n = 19, 11, 19, 11, 20, 11, 6, 6 zygotes. (F) CHIN-1 is essential for the cortical polarization of PAR-6 in zygotes where PAR-2 is predominant. Representative images of mCherry::PAR-6 and GFP::PAR-2 in zygotes under control, *par-6(pRNAi)*, *par-6(pRNAi);chin-1(RNAi)*, and *chin-1(RNAi)* conditions are shown. *chin-1(RNAi)* caused depolarization of mCherry::PAR-6, resulting in the weak enrichment of mCherry::PAR-6 throughout the cortex. Scale Bar, 5 μm. (G) The graph shows the size of cortical mCherry::PAR-6 domain in zygotes under conditions shown in (F). Data present mean ± s.d. from n = 16, 17, 30, 6 zygotes. (B-E and G) *p*-values; Mann–Whitney test.

The above results suggest that the polarized cortical PAR domains can be maintained by an inter-connected network involving two reciprocal exclusion pathways (Figure 4A). One pathway depends on the interactions among PAR-3–PKC-3–PAR-6 at the anterior cortex and between PAR-1–PAR-2 at the posterior cortex. In this pathway, antagonistic phosphorylation between the two complexes contributes to reciprocal cortical exclusion. The other pathway relies on interactions among CDC-42–PAR-6–PKC-3 at the anterior cortex and CHIN-1 at the posterior cortex. Here PKC-3 excludes CHIN-1 from the anterior cortex via an unidentified mechanism, whereas CHIN-1 inactivates and excludes CDC-42 from the posterior cortex. In contrast to our revised model, the conventional model of cortical PAR network includes only one reciprocal exclusion pathway involving PAR-3–PKC-3–PAR-6–CDC-42 at the anterior cortex and PAR-1–PAR-2 at the posterior cortex (Blanchoud et al., 2015; Dawes and Munro, 2011; Goehring et al., 2011; Gross et al., 2019). A recent study proposed an inter-connected network comprising PAR-3–PKC-3–PAR-6–CDC-42 at the anterior cortex and PAR-1–PAR-2 and CHIN-1 at the posterior cortex (Sailer et al., 2015). However, the earlier models assumed that PAR-3 is essential for the local recruitment and maintenance of PAR-6 at the anterior cortex (Blanchoud et al., 2015; Dawes and Munro, 2011; Goehring et al., 2011; Gross et al., 2019; Sailer et al., 2015), and therefore cannot explain the maintenance of the polarized PAR-6 domain even in the absence of cortical PAR-3 loading.

**Figure 4.**
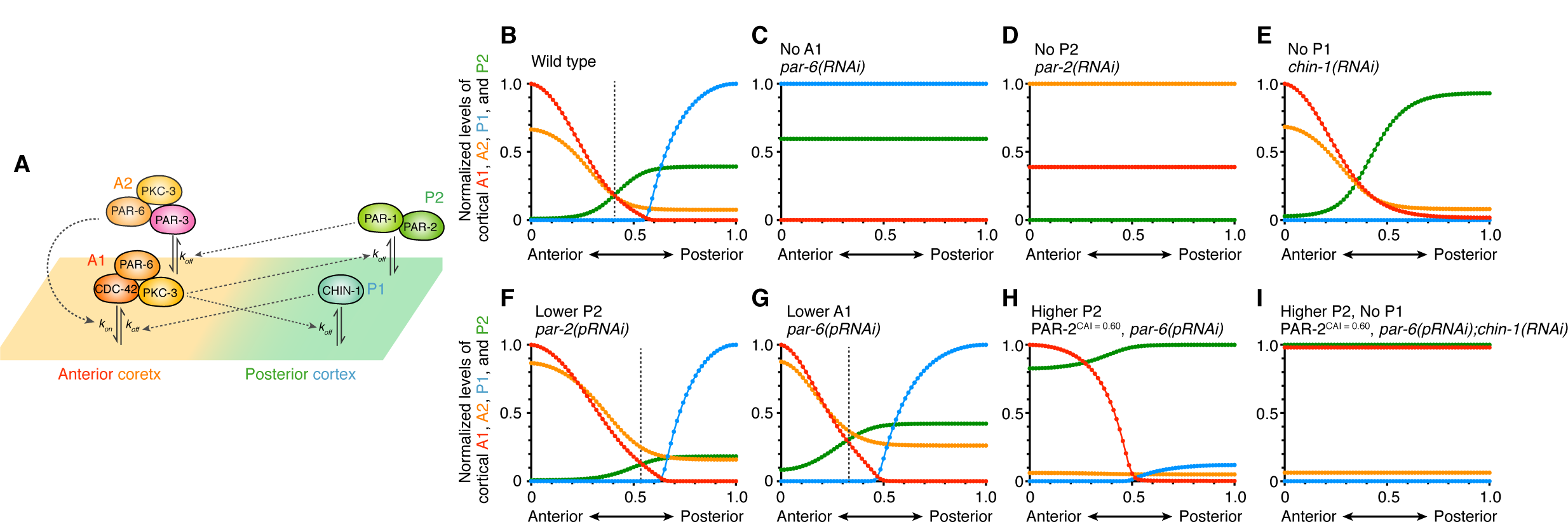
Modelling of a combinatorial network of two reciprocal cortical-exclusion pathways. (A) Schematic view of a model of cortical PAR polarity patterning. A mutually-exclusive pathway between A_2_ (PAR-3, PAR-6, and PKC-3) and P_2_ (PAR-1 and PAR-2) is complemented by another mutually-exclusive pathway between A_1_ (CDC-42, PAR-6, and PKC-3) and P_1_ (CHIN-1). This scheme includes four inhibitory interactions (from A_1_ to P2, from P_2_ to A2, from A_1_ to P1, and from P_1_ to A1) and one positive interaction (from A_2_ to A1). (B-I) Steady-state analysis of the model equations in zygotes under following conditions. Predicted distributions of four PAR species at the cortex along the anteroposterior axis are shown. (B) wild-type, (C) No A_1_ (*par-6(RNAi)*), (D) No P_2_ (*par-2(pRNAi)*), (E) No P_1_ (*par-6(pRNAi)*), (F) Lower P_2_ (*par-2(pRNAi)*), (G) Lower A_1_ (*par-6(pRNAi)*), (H) Higher P_2_ (GFP::PAR-2^CAI = 0.60^, *par-6(pRNAi)*), (I) Higher P_2_ and No P_1_ (GFP::PAR-2^CAI = 0.60^, *par-6(pRNAi); chin-1(RNAi)*).

### Simulation of PAR patterning in zygotes

To test if the inter-connected PAR network involving two reciprocal exclusion pathways would sufficiently explain the PAR patterning in zygotes with manipulated PAR balances, we developed a mathematical model of PAR patterning network. Previous studies simulated only a sub-circuit involving one reciprocal cortical-exclusion pathway either between PAR-3–PKC-3–PAR-6 and PAR-1–PAR-2 (Blanchoud et al., 2015; Dawes and Munro, 2011; Goehring et al., 2011) or between PKC-3–PAR-6–CDC-42 and CHIN-1 (Sailer et al., 2015). We developed a model in which the anterior cortical domain contains two species, A_1_ [CDC-42–PAR-6–PKC-3] and A_2_ [PAR-3–PAR-6–PKC-3], and the posterior cortical domain includes two species, P_1_ [CHIN-1] and P_2_ [PAR-1–PAR-2] (Figure 4A). We included four inhibitory interactions: from A_1_ [CDC-42–PAR-6–PKC-3] to P_2_ [PAR-1–PAR-2], from P_2_ [PAR-1–PAR-2] to A_2_ [PAR-3–PAR-6– PKC-3], from A_1_ [CDC-42–PAR-6–PKC-3] to P_1_ [CHIN-1], and from P_1_ [CHIN-1] to A_1_ [CDC-42–PAR-6– PKC-3] (Figure 4A). As PAR-3 facilitates the cortical recruitment of PAR-6–PKC-3 during the early stage of polarization, we also included a positive interaction from A_2_ [PAR-3–PAR-6–PKC-3] to A_1_ [CDC-42– PAR-6–PKC-3] (Figure 4A). A steady-state analysis of the four PAR species in a wild-type condition revealed the enrichment of A_1_ [CDC-42–PAR-6–PKC-3] and A_2_ [PAR-3–PAR-6–PKC-3] at the anterior cortex and that of P_1_ [CHIN-1] and P_2_ [PAR-1–PAR-2] at the posterior cortex (Figure 4B). Subsequent simulation wherein the concentration of A_1_ [CDC-42–PAR-6–PKC-3] (Figure 4C), P_2_ [PAR-1–PAR-2] (Figure 4D), or P_1_ [CHIN-1] (Figure 4E) were reduced to zero reproduced the PAR patterns observed in zygotes where either PAR-6, PAR-2, or CHIN-1 had been depleted by RNAi (Figures 4C-4E). Therefore, this model confirms the essential roles of PAR-2 and PAR-6 and a non-essential role of CHIN-1 in a wild-type background. A reduction in the total concentration of P_2_ [PAR-1–PAR-2] reduced the size of the P_2_ [PAR-1–PAR-2] domain and shifted the anterior-posterior boundary to the posterior pole (Figure 4F). In contrast, a decrease in the total concentration of A_1_ [CDC-42–PAR-6–PKC-3] enlarged the size of the P_2_ [PAR-1–PAR-2] domain and shifted the anterior-posterior boundary to the anterior pole (Figure 4G). Further overexpression of P_2_ [PAR-1–PAR-2] induced a bifurcation characterized by high and low cortical concentration of P_2_ [PAR-1–PAR-2] and A_2_ [PAR-3–PAR-6–PKC-3], respectively, throughout the cortex, while maintaining an anteriorly-enriched A_1_ [CDC-42–PAR-6–PKC-3] domain (Figure 4H). At the bifurcation point where P_2_ [PAR-1–PAR-2] localizes throughout the cortex, a reduction in P_1_ [CHIN-1] led to depolarization of the anterior A_1_ [CDC-42–PAR-6–PKC-3] domain (Figure 4I), indicating the requirement for CHIN-1 to maintain a polarized PAR-6 domain. Thus, our simulation based on the inter-connected network involving two reciprocal exclusion pathways could recapitulate the PAR patterning in zygotes subjected to manipulation of the balance between PAR proteins. These analyses reveal that the combinatorial network maintains the polarization of PAR-6 independently of reciprocal exclusion between PAR-2 and PAR-6 in zygotes.

### Reciprocal exclusion of PAR proteins is not essential for unequal inheritance of fate determinants

We next investigated whether a state maintaining an anteriorly-polarized PAR-6 domain in the absence of reciprocal exclusion between PAR-2 and PAR-6 would yield unequal inheritance of fate determinants during the first cell division. The zygotes with manipulated PAR balances were categorized into three classes based on the distribution patterns of both GFP::PAR-2 and cytoplasmic factors including the fate determinants (P-granules and MEX-5) and the position of a cleavage furrow (specified by the position of the mitotic spindle) (see Figure 5 legend for the definition of the three phenotypic classes). More than half of the zygotes with a predominance of PAR-2 (GFP::PAR-2^CAI = 0.60^; mCherry::PAR-6, *par-6(pRNAi)* for 9 hours) localized GFP::PAR-2 throughout the cortex but exhibited the accumulation of PGL-1-positive P-granules within the posterior cytoplasm (class II zygotes in Figures 5A, 5B, and 5E). Although these zygotes contained fewer P-granules than wild-type zygotes, these granules were polarized toward the posterior cortex when compared to zygotes that severely depleted of PAR-6 (GFP::PAR-2CAI = 0.60; mCherry::PAR-6, *par-6(RNAi)* for 24 hours) (Figure 5A). These PAR-2-predominant zygotes also established a gradient of MEX-5 enriched within the anterior cytoplasm (class II zygotes in Figures 5A, 5C, and 5F). The mitotic spindles were displaced toward the posterior cortex, resulting in a cleavage furrow positioned toward the posterior pole (class II zygotes in Figures 5A, 5D, and 5G). Both manipulated and control embryos retained cell-cycle asynchrony between AB and P1 cells in two-cell stage embryos (Figures 6A and S6A). These observations suggest that these asymmetries between AB and P1 cells were established independently of reciprocal cortical exclusion between PAR-2 and PAR-6 during the first cell division.

**Figure 5.**
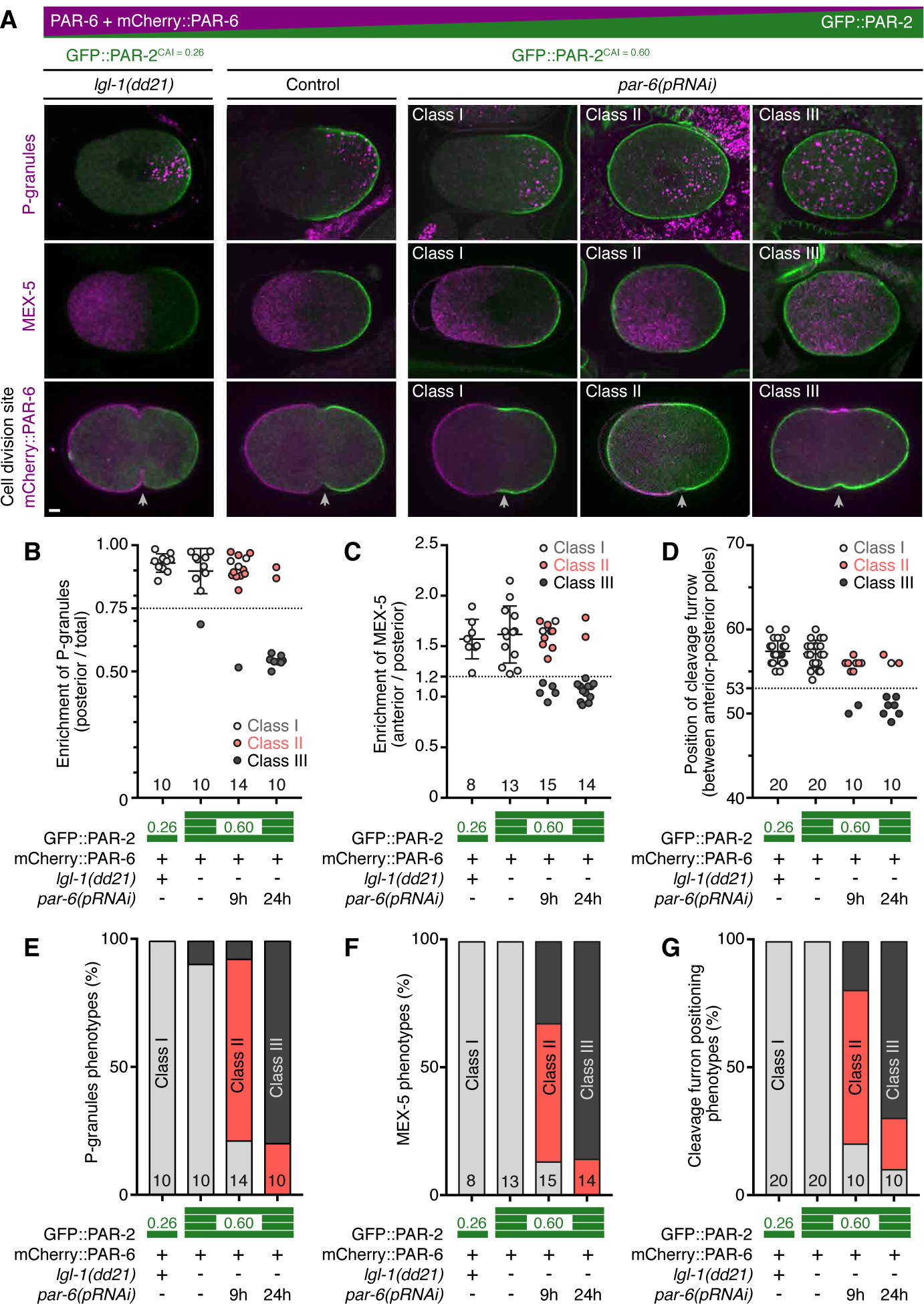
Fate determinants can be segregated independently of reciprocal cortical exclusion between PAR proteins. (A) Representative images of the distribution of GFP::PAR-2 (green) and the cytoplasmic factors including P-granules (magenta), MEX-5 (magenta), and the position of a cleavage furrow (white arrowheads) in zygotes where the balance between PAR-6 and PAR-2 were manipulated. These zygotes were classified into three groups: Class I zygotes established the polarized distributions of GFP::PAR-2 and the cytoplasmic factors. Class II zygotes exhibited the polarized distribution of the cytoplasmic factors, whereas GFP::PAR-2 localized throughout the cortex. Class III zygotes showed uniform distributions of GFP::PAR-2 and the cytoplasmic factors. Scale Bar, 5 μm. (B-D) The graphs depict the enrichment of P-granules (B) and MEX-5 (C) in the anterior and the posterior cytoplasm, and the position of cleavage furrows (D) in zygotes described in (A). (E-G) The graphs depict the percentage of three phenotypic classes shown in (B-D). (B-G) The number of samples are indicated in the graphs. Height of green bars indicate the CAI values of *gfp::par-2* transgene.

**Figure 6.**
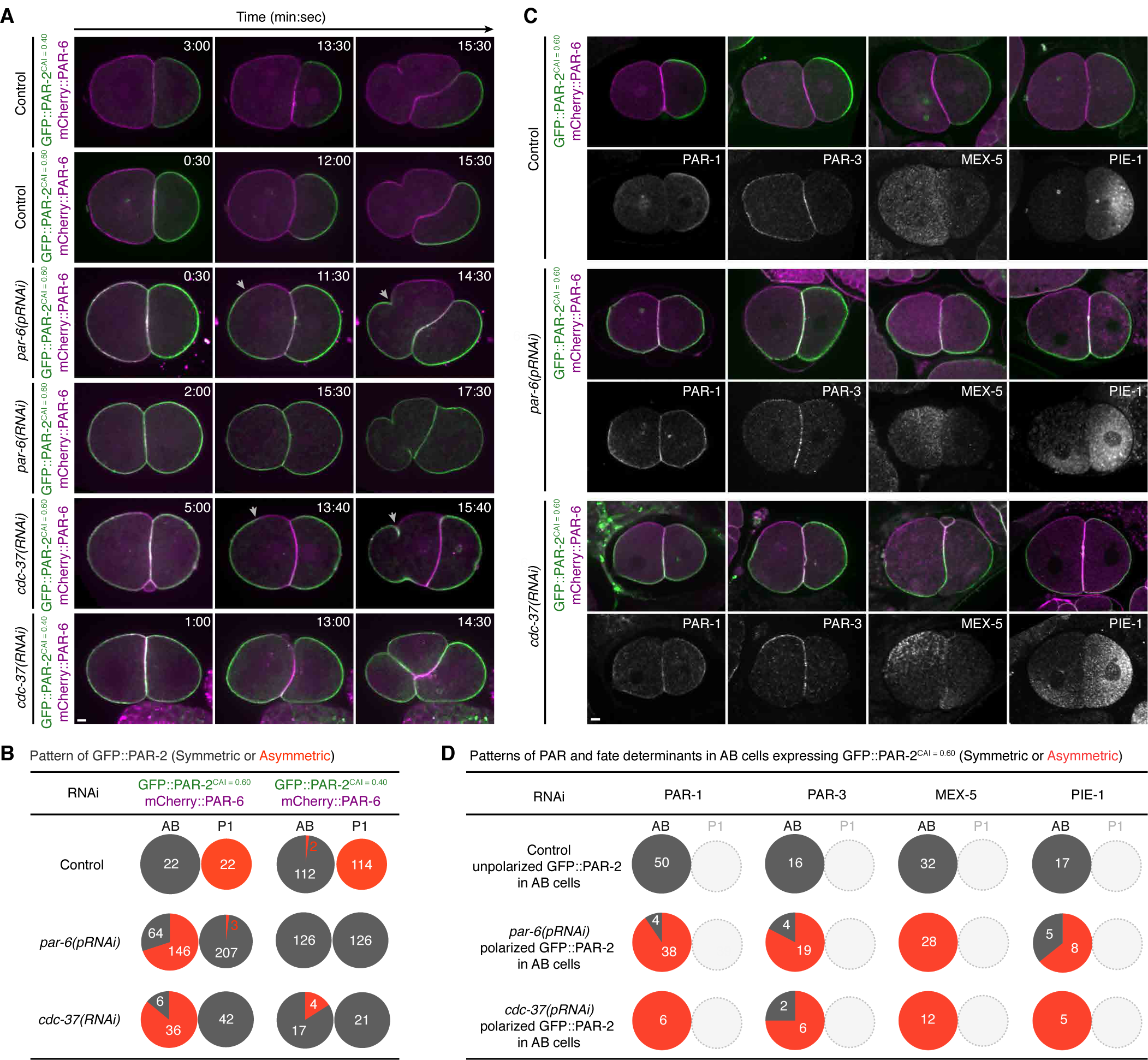
The balance between PAR proteins defines the choice between the two division modes. (A) Changes in the levels of PAR proteins are sufficient to reverse the choice between the two cell division modes in two-cell stage embryos. Representative time-lapse images of embryos expressing mCherry::PAR-6 (magenta) and either GFP::PAR-2CAI = 0.40 or GFP::PAR-2CAI = 0.60 (green) under control*, par-6(pRNAi), par-6(RNAi)*, and *cdc-37(RNAi)* conditions are shown. Arrowheads show the boundary between mCherry::PAR-6 and GFP::PAR-2 cortical domains in AB cells. The times stated are with respect to the completion of cytokinesis in zygotes. Scale Bar, 5 μm. (B) The graphs depict the percentage of AB and P_1_ cells that underwent either equal or unequal inheritance of GFP::PAR-2 under control*, par-6(pRNAi)*, and *cdc-37(RNAi)* conditions. (C) Induced asymmetry in cortical PAR proteins mediates the segregation of fate determinants in AB cells. Representative images of the distributions of PAR-1, PAR-3, MEX-5, and PIE-1 in embryos expressing both mCherry::PAR-6 (magenta) and GFP::PAR-2CAI = 0.60 (green) under control, *par-6(pRNAi)*, and *cdc-37(RNAi)* conditions are shown. Scale Bar, 5 μm. (D) The graphs depict the percentage of AB cells that segregated PAR-1, PAR-3, MEX-5, and PIE-1 between their daughter cells. (B and D) The number of cells observed is indicated in the graphs.

### The balance between antagonizing PAR proteins defines the binary specification of the division modes in two-cell stage embryos

Genetic manipulation of the PAR balances enabled us to establish the PAR-2-predominant zygotes, wherein the cortical pattern of PAR proteins was uncoupled from the segregation of fate determinants and the mitotic spindle displacement during the first cell division. Therefore, we were able to test whether these manipulations in the PAR protein levels would affect the specification of the division modes in AB and P1 cells, while ensuring the correct asymmetries in fate determinants, cell size, and cell-cycle progression between these sister cells. The invariant lineage of *C. elegans* dictates the commitment of AB and P1 cells to the symmetric and asymmetric modes of cell division, respectively. In control embryos expressing normal levels of PAR-6 and PAR-2, the AB cells localized mCherry::PAR-6 and PAR-3 throughout the cortex and GFP::PAR-2 and MEX-5 within the cytoplasm, leading to equal inheritance of PAR proteins (GFP::PAR-2, mCherry::PAR-6, and PAR-3) and fate determinants (P-granules, MEX-5, and PIE-1) between the two daughter cells (Figures 6A-6D). PAR-2-predominant AB cells exhibited overlapping distributions of mCherry::PAR-6 and GFP::PAR-2 throughout the cortex during early prophase (Figure 6A). Subsequently, mCherry::PAR-6 and GFP::PAR-2 began to polarize into two distinct cortical domains, resulting in unequal inheritance of GFP::PAR-2 and mCherry::PAR-6 between the two daughter cells (Figures 6A, 6B, and S6A). The orientation of the PAR-6 and PAR-2 cortical domains occurred randomly with respect to the contact between AB and P1 (Figure S6C), suggesting that the PAR patterning is not instructed by the cell-cell contact. The polarization pattern of GFP::PAR-2 and mCherry::PAR-6 in AB cells was strongly associated with the polarization of PAR-1 and PAR-3 to their respective cortical domains. Remarkably, the PAR-2-predominant AB cells partitioned cytoplasmic MEX-5 and PIE-1 toward the mCherry::PAR-6 domain and the GFP::PAR-2 domains, respectively (Figures 6C, 6D, and S7), indicating that the polarized PAR domains are able to induce the asymmetric mode of AB cell division. During anaphase, mitotic spindle rotation was not detectable in AB cells in which both GFP::PAR-2 and mCherry::PAR-6 were polarized (Figures 6A and S6C). In contrast to AB cells, P1 cells from the same embryos exhibited uniformly distributed GFP::PAR-2 at the cortex and mCherry::PAR-6 in the cytoplasm, resulting in the symmetric mode of P1 cell division (Figures 6A-6C, S6, and S7). We thus conclude that when other relevant factors are controlled, the manipulation of PAR protein levels is sufficient to reverse the specification of the division modes between AB and P1 cells.

To further test whether changes in the balance of PAR proteins could sufficiently induce asymmetric mode of AB cell division, we characterized other mutant embryos in which PAR-1 and PAR-2 had been inherited equally between AB and P1 cells. In a zygote, mutual exclusion between the anterior and the posterior PAR domains requires CDC-37 (Beers and Kemphues, 2006). Here, *cdc-37(RNAi)* treatment induced overlapping cortical distributions of mCherry::PAR-6 and GFP::PAR-2, and the symmetric distributions of MEX-5 and P-granules during the first cell division (Figures S4A and S4C-S4E). Notably, higher levels of GFP::PAR-2^CAI = 0.60^, but not moderate levels of GFP::PAR-2^CAI = 0.40^, induced the asymmetric segregation of GFP::PAR-2, PAR-1, PAR-3, MEX-5, and PIE-1 in AB cells of two-cell stage *cdc-37(RNAi)* embryos (Figures 6A-6D, S6A, and S7). These results further confirmed that a change in the PAR protein balance can sufficiently reverse the selection of the division modes between AB and P1 cells.

Next, we investigated whether an optimum level of PAR proteins would enable the asymmetric mode of cell division in both sister cells that had equally inherited fate determinants during the first cell division. Here we aimed to verify whether the specification of the division modes in AB and P1 cells was truly independent of the unequal inheritance of fate determinants and asymmetries in cell size and cell-cycle progression. Using RNAi to deplete cyclin-E (CYE-1) that prevented the polarization of PAR proteins via a failure of centrosome-mediated polarization in zygotes (Cowan and Hyman, 2006), we observed the equal inheritance of PAR proteins (mCherry::PAR-6, GFP::PAR-2, and PAR-3) and fate determinants (MEX-5, PIE-1, and P-granules) during the first cell division (Figures S5B, S5C, S5E, and S5F). Moreover, both daughter cells exhibited equal sizes and synchronized cell-cycle progression (Figures 7A and S6B). In two-cell stage *cye-1(RNAi)* embryos expressing lower levels of GFP::PAR-2^CAI = 0.26^, both cells equally inherited GFP::PAR-2 during mitosis (Figures 7A and 7B). Remarkably, in two-cell stage *cye-1(RNAi)* embryos expressing higher levels of GFP::PAR-2CAI = 0.40, both cells segregated GFP::PAR-2 to the polar cortex. Consequently, GFP::PAR-2 was inherited unequally by one daughter cell during the second cell division (Figures 7A, 7B, and S6B). The polarized distribution of PAR-2 was strongly associated with the segregation of PAR-3 into the respective cortical domain, of MEX-5 in the cytoplasm opposite to the GFP::PAR-2 domain, and of PIE-1 and P-granules toward the GFP::PAR-2 domain (Figures 7C, 7D, and S7). The polarized distribution of PAR proteins in two-cell stage embryos was often associated with the rotation of mitotic spindles toward the cell-cell contact site (Figure S6C). Therefore, we conclude that the selected mode of cell division in two-cell stage embryos is completely independent of the unequal inheritance of fate determinants, cell-size asymmetry and cell-cycle asynchrony between the sister cells. Our findings support a model that the balance between antagonizing PAR proteins is the critical intrinsic cue for binary specification regarding the asymmetric or symmetric mode of cell division.

**Figure 7.**
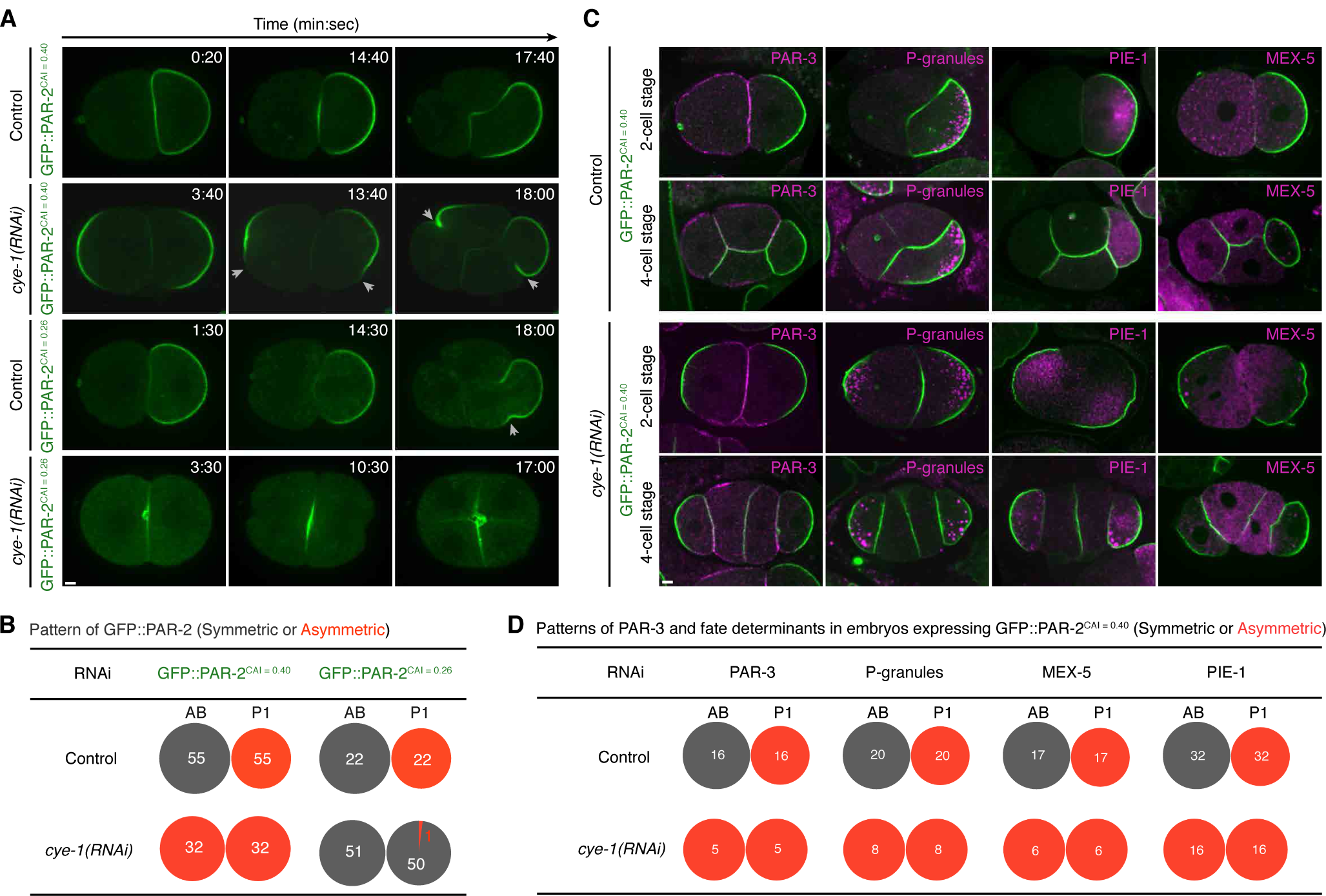
The choice between the two division modes is independent of asymmetries in fate determinants between the sister cells. (A) Unequal inheritance of fate determinants and asymmetries in cell size and cell-cycle progression are not required for determining the two modes of cell division. Representative time-lapse images of embryos expressing either GFP::PAR-2^CAI = 0.40^ or GFP::PAR-2^CAI = 0.26^ (green) under control and *cye-1(RNAi)* conditions are shown. Arrowheads show the edges of cortical GFP::PAR-2 domains. The times stated are with respect to the completion of cytokinesis in zygotes. Scale Bar, 5 μm. (B) The graphs depict the percentage of AB and P_1_ cells that underwent either equal or unequal inheritance of GFP::PAR-2 between their daughter cells. (C) Induced asymmetry in cortical PAR proteins in two-cell stage *cye-1(RNAi)* embryos mediates unequal inheritance of fate determinants. Representative images of the distributions of PAR-3, PGL-1-positive P-granules, PIE-1, and MEX-5 (magenta) in embryos expressing GFP::PAR-2CAI = 0.40 (green) under control and *cye-1(RNAi)* conditions are shown. Scale Bar, 5 μm. (D) The graphs depict the percentage of the sister cells in two-cell stage embryos that segregated PAR-3, P-granules, PIE-1, and MEX-5 during the second cell division. (B and D) The number of cells observed is indicated in the graphs.

### Simulation of PAR patterning in two-cell stage embryos

To test if the binary specification of the division mode in two-cell stage embryos can be explained by unequal inheritance of the balance between antagonizing PAR proteins, we applied a mathematical model of the aforementioned PAR network to two-cell stage embryos (Figures 8A-8E). We performed steady-state analysis of the four PAR species distribution in AB and P1 cells, which are derived from wild-type zygotes or the zygotes with a predominance of PAR-2 (Class II zygotes in Figure 5). Inheritance of the PAR species between AB and P1 cells and their initial distribution in AB and P1 cells were defined by their steady-state distribution in zygotes (Figures 4 and 8A). These PAR species were partitioned at the position of cleavage furrow measured in zygotes (56.1 ± 1.6 % in wild-type zygotes, and 54.8 ± 2.3% in PAR-2-predominant zygotes) (Figures 5 and 8A).

**Figure 8.**
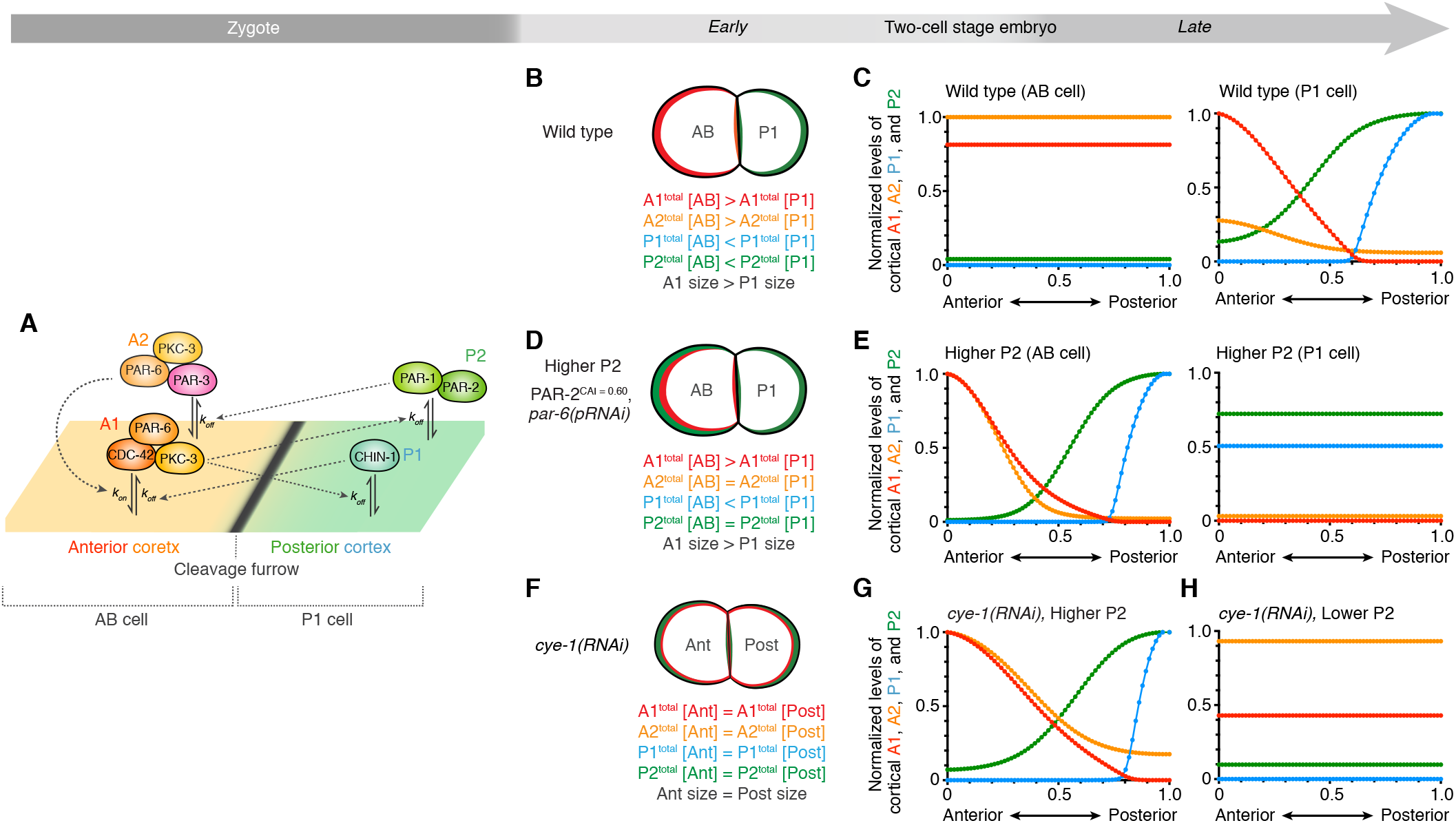
Modelling of a combinatorial PAR network in two-cell stage embryos. (A) Schematic view of PAR proteins inheritance in a wild-type embryo. The four PAR species at the steady state in a zygote are partitioned at the cleavage furrow, resulting in unequal inheritance of these PAR species into AB and P_1_ cells. (B and C) Steady-state analysis of the PAR network model in two-cell stage wild-type embryo. (B) Schematic of PAR-6 (red) and PAR-2 (green) distribution in an AB cell and a P_1_ cell. (C) Predicted distributions of the PAR species at the cortex along the anteroposterior axis in an AB cell (left) and a P_1_ cell (right). (D and E) Steady-state analysis of the PAR network model in two-cell stage embryo with a predominance of PAR-2. (D) Schematic of PAR-6 (red) and PAR-2 (green) distribution in an AB cell and a P_1_ cell. (E) Predicted distributions of the PAR species at the cortex along the anteroposterior axis in an AB cell (left) and a P_1_ cell (right). (C and E) Cortical concentration of each PAR species is normalized between AB and P_1_ cells in the same embryo. (F-H) Steady-state analysis of the PAR network model in two-cell stage *cye-1(RNAi)* embryo. (F) Schematic of PAR-6 (red) and PAR-2 (green) distribution in a *cye-1(RNAi)* embryo. (G and H) Predicted distributions of the PAR species at the cortex along the anteroposterior axis in the daughter cells expressing higher levels of PAR-2 (G) or lower level of PAR-2 (H). Cortical concentration of each PAR species is normalized between these two conditions.

Wild-type zygotes partitioned the PAR species unequally into two daughter cells. AB cells inherited higher concentration of A_1_ [CDC-42–PAR-6–PKC-3] and A_2_ [PAR-3–PAR-6–PKC-3] and lower concentration of P_1_ [CHIN-1] and P_2_ [PAR-1–PAR-2] (Figure 8B). The total concentration of the PAR species in P1 cells was determined according to the law of mass conservation (Figure 8B). In wild-type two-cell stage embryos, AB cells exhibited symmetric distribution of all PAR species, while P1 cells segregated them into two distinct cortical domains (Figure 8C). In contrast to wild-type zygotes, PAR-2-predominant zygotes underwent almost-equal inheritance in A_2_ [PAR-3–PAR-6–PKC-3] and P_2_ [PAR-1– PAR-2], while keeping unequal inheritance in A_1_ [CDC-42–PAR-6–PKC-3] and P_1_ [CHIN-1] into two daughter cells (Figure 8D). In PAR-2-predominant two-cell stage embryos, AB cells segregated A_1_ [CDC-42–PAR-6–PKC-3] and A_2_ [PAR-3–PAR-6–PKC-3] at the anterior cortex and P_1_ [CHIN-1] and P_2_ [PAR-1– PAR-2] at the posterior cortex (Figure 8E), while P_1_ cells exhibited symmetric distribution of all PAR species (Figure 8E). The PAR patterning in the PAR-2-predominant embryos relied on unequal inheritance in A_1_ [CDC-42–PAR-6–PKC-3], but not in P_1_ [CHIN-1], between AB and P1 cells (Figure S8).

We also performed steady-state analysis of the PAR species distribution in two-cell stage *cye-1(RNAi)* embryos that expressed different levels of PAR-2 (Figures 8F and 8G). *cye-1(RNAi)* zygotes partitioned all PAR species equally and divided symmetrically in size, making two identical daughter cells (Figures S5 and 8F). Because the mechanism triggering symmetry breaking in *cye-1(RNAi)* embryos is unknown, we analysed a steady-state solution from an initial condition with transient random fluctuation of all four PAR species at the cortex. When the level of P_2_ [PAR-1–PAR-2] was higher (up to 2.5-fold increase compared to the wild-type condition), both daughter cells segregated A_1_ [CDC-42–PAR-6–PKC-3] and A_2_ [PAR-3–PAR-6–PKC-3] at the anterior cortex and P_1_ [CHIN-1] and P_2_ [PAR-1–PAR-2] at the posterior cortex (Figure 8G). Reduction of the P_2_ [PAR-1–PAR-2] level (to less than 50%) caused symmetric distribution in all PAR species in both daughter cells (Figure 8H), indicating the P_2_ [PAR-1–PAR-2] level as one of the bifurcation parameters that switch PAR patterning between apolar and polar states. Therefore, our simulation could recapitulate the PAR patterning in two-cell stage embryos subjected to manipulation of the balance between PAR proteins. These analyses reveal that unequal inheritance of the balance between antagonizing PAR proteins plays a critical role in the binary specification of PAR patterning in two-cell stage embryos.

## DISCUSSION

In this report, we presented evidence indicating that the binary specification between asymmetric and symmetric modes of cell division relies on the balance between antagonizing PAR proteins in *C. elegans* embryos. We demonstrated that manipulating the levels of two PAR proteins (PAR-2 and PAR-6) inherited during the first cell division could induce all combinations of asymmetric and/or symmetric pattern of PAR proteins in their daughter cells. The polarized PAR domains artificially induced in these daughter cells were sufficient to establish the unequal inheritance of fate determinants, resulting in successful induction of asymmetric cell division. These results indicate that changes in the balance between PAR proteins are sufficient to re-program the otherwise invariable lineage of cell division modes. Moreover, the division mode adopted occurred independently of the unequal inheritance of fate determinants, cell-size asymmetry, and cell-cycle asynchrony between the two sister cells. Given that the division modes in two-cell stage embryos are directed in a cell-autonomous manner (Priess and Thomson, 1987), the intrinsic cue that specifies the division mode must be unequally inherited during the first cell division. We propose that the balance between antagonizing PAR proteins, which should be inherited unequally during the first cell division, is the previously-unknown intrinsic cue used to specify the division modes in AB and P1 cells. Our simulation also support that the balance between antagonizing PAR proteins can be a bifurcation parameter that controls the self-organizing interactions among PAR proteins in two-cell stage embryos. A certain balance of PAR protein levels promotes the self-organization of antagonizing PAR proteins into a mutually-exclusive pattern at the cortex, which then mediates the asymmetric segregation of fate determinants. In contrast, a distinct balance that does not permit the self-organizing interactions results in the maintenance of unpolarized PAR domains, which causes the equal segregation of fate determinants. During normal development, P1 cells may use such self-organizing mechanism to re-establish the asymmetric distribution of PAR proteins, whereas AB cells exhibit a disproportion in PAR proteins that blocks the self-organization.

PAR proteins have been generally considered as passive effectors of the intrinsic cell-division program. Especially, the segregation of PAR proteins is induced by the intrinsic cue from centrosomes in *C. elegans* zygotes (Klinkert et al., 2019; Reich et al., 2019; Zhao et al., 2019). In contrast, our findings highlight that the self-organizing property of antagonizing PAR proteins serves as an active cue that instructs the intrinsic cell-division program in two-cell stage embryos. Recent studies observed delayed polarization of PAR proteins in zygotes depleted of a centrosome-mediated cue, Aurora-A (Klinkert et al., 2019; Reich et al., 2019; Zhao et al., 2019), suggesting the centrosome-independent self-organizing property of PAR proteins. We propose that unequal inheritance of the self-organizing property of PAR proteins plays a critical role in the binary specification of PAR patterning in two-cell stage embryos. It will be necessary to test if manipulating the PAR balance in other cell types can be sufficient to induce self-organization of PAR proteins and thus promote asymmetric mode of cell division during later stages of development.

Conventionally, the PAR network involves a two-node topology comprising a single reciprocal exclusion pathway among the PAR proteins (Blanchoud et al., 2015; Chau et al., 2012; Dawes and Munro, 2011; Goehring et al., 2011; Gross et al., 2019; Hoege and Hyman, 2013). A recent study proposed a cross-inhibitory network between PAR-3–PKC-3–PAR-6–CDC-42 at the anterior cortex and PAR-1–PAR-2 and CHIN-1 at the posterior cortex (Sailer et al., 2015). However, this model assumes that PAR-3 is required for the cortical recruitment of PAR-6 in the steady state. Here, we demonstrated that a simple two-node pathway of PAR proteins (one containing PAR-3–PAR-6–PKC-3 and the other, PAR-1–PAR-2) is complemented by another pathway of reciprocal cortical exclusion. This secondary pathway involves the polarized activity of CDC-42 (one node containing CDC-42–PAR-6–PKC-3 and the other, CHIN-1). Our simulations based on a combinatorial network of two reciprocal exclusion pathways recapitulate the cortical patterning of PAR proteins in zygotes under various balances of PAR proteins. Our observations of *C. elegans* zygotes also highlight that this combinatorial network ensures the cortical polarization of PAR-6 even in the absence of (1) cortical enrichment of PAR-3 and (2) reciprocal exclusion between PAR-2 and PAR-6. Indeed, the robustness of the combinatorial network enables uncoupling the inheritance of PAR proteins from that of fate determinants during the first cell division. These findings are consistent with previous observations wherein several alleles of *par* mutants failed to establish the reciprocal exclusion of PAR proteins but induced the polarization of P-granules within the cytoplasm (Bowerman et al., 1997; Boyd et al., 1996; Kemphues et al., 1988; Morton et al., 2002). Our results further reveal that the reciprocal exclusion of PAR proteins in zygotes is not essential for the unequal inheritance of fate determinants but is necessary for the maintenance of invariable cell-division lineage in two-cell stage embryos.

Previous studies have implicated PAR polarity in cortical polarization and asymmetric cell division in many cell types, including *Drosophila* neuroblasts (Knoblich, 2010), vertebrate embryos (Maitre et al., 2016), neural progenitor cells (Homem et al., 2015), and several types of stem cells (Chang et al., 2007; Dumont et al., 2015; Lechler and Fuchs, 2005). The striking conservation of the function of PAR proteins in both invertebrates and vertebrates raises a possibility that the self-organizing PAR system is a conserved executor of the binary decision between the two division modes. Interestingly, the ratios of PAR proteins can be modified dynamically during development. For example, the ratio of PAR-3 to PAR-1 is actively modified to induce morphogenetic movement in *Drosophila* epithelium (Wang et al., 2012). It is therefore tempting to speculate that the balance between antagonizing PAR proteins could be controlled dynamically to affect the facultative choice between the two modes of cell division during development. Indeed, many types of stem cells shift between these modes to balance the needs of self-renewal and cell differentiation (Morrison and Kimble, 2006). Hence, our findings suggest that the dynamic control of cortical polarization is directly linked to the binary specification regarding the asymmetric or symmetric mode of cell division, a critical decision faced by every single cell in a multicellular organism.

## ACKNOWLEDGMENTS

This study was supported by the Singapore National Research Foundation (NRF-NRFF2012-08 [F.M.]), and the Strategic Japan-Singapore Cooperative Research Program by the Japan Science and Technology Agency and the Singapore Agency for Science, Technology, and Research (1514324022 [F.M.] and 15658064 [T.S.]). We are grateful to Nathan Goehring (The Francis Crick Institute), Pierre Gonczy (EPFL), Monica Gotta (University of Geneva), Carsten Hoege and Anthony Hyman (MPI), Ken Kemphues (Cornel University), Jean-Claude Labbe (IRIC), Shibi Mathew (Temasek Life-sciences Laboratory), Geraldine Seydoux (Johns Hopkins University), Shigeo Ohno (Yokohama-City University), Zhen Zhang and Pakorn Kanchanawong (Mechanobiology Institute, Singapore), and the Caenorhabditis Genetic Center for strains, reagents and expertise. We also thank Andrew Wong (Mechanobiology Institute, Singapore) and members in the Motegi lab for comments on the manuscript.

## AUTHOR CONTRIBUTIONS

The experimental design and presented ideas were developed together by all authors. F.M. guided the study and wrote the manuscript with input from all authors. Y.W.L. performed all experiments. F.L.W., P.S., and T.S. developed the theoretical models for an inter-connecting network of two reciprocal cortical exclusion pathways.

## METHODS

### *C. elegans* strains and RNAi

*C. elegans* transgenes were constructed in the Gateway destination vector (pID3.01 or pID2.02) and transformed into worms by biolistic transformation. The strains used in this study are listed in Table S1. Strains were maintained at 20°C and shifted to 25°C for 20–30 hours before recording. RNAi experiments, except for *par-6(3’UTR RNAi)*, were performed by the feeding method. The RNAi constructs used in this study are listed in Table S2. L4440-based RNAi clones were transformed into *Escherichia coli* (*E. coli*) HT115 cells. The transformants were grown at 37°C in liquid LB media supplemented with 500 µg/mL carbenicillin. A volume of 100 µL of transformed *E.coli* liquid culture was seeded onto Nematode Growth Media (NGM) plates with 1 mM IPTG, and incubated at room temperature overnight. Worms at the L3/L4 stage were transferred to feeding RNAi plates and incubated at 25°C for 24 hours. *par-6(3’UTR RNAi)* was performed by the soaking method. *par-6* 3’ UTR sequence was prepared by PCR with following primers and genomic DNA of N2 worms as a template.

5’-GTAATACGACTCACTATAGGGC aaaactcttttcagccatttttcc–3’

5'-GCGTAATACGACTCACTATAGGGC tcactaataatgtgaatttcagg-3’

Double-stranded RNA (dsRNA) was prepared by *in vitro* transcription of a PCR product with T7 RNA polymerase. Worms at the L4 stage were incubated in a solution containing dsRNA at concentration 1 mg ml^-1^ at 20 ºC for 24 hours, and then grown on an NGM plate at 25ºC. Their phenotypes were observed 24 hours after removal of from the dsRNA solution.

### Imaging of *C. elegans*

For live imaging, embryos were isolated from gravid hermaphrodite animals into egg salt buffer, placed on coverslips, and inverted on to slides with 20 μm monodisperse polysterene beads (Bangs Laboratories, Inc). Embryos were observed at 25°C with a CFI Plan Apochromat 60× N.A.1.4 oil immersion lens on a Nikon Ni-E motorized upright microscope (Nikon) fitted with a CSU-X1 spinning disk confocal system (Yokogawa Electric Corp.) with LaserStack 491, 561, and 642 solid-state diode lasers (Intelligent Imaging Innovation Inc.). Images were acquired with a Photometrics Evolve512 camera (Photometrics) controlled by Metamorph software (Intelligent Imaging Innovation Inc.) using a 250 ms exposure at 20% power on the 491 and 561 lasers and 1×1 binning in the camera. Nuclear envelope breakdown (NEBD) was defined as the first frame when the GFP fusion was no longer excluded from pronuclei.

For immunofluorescence staining, embryos were isolated into egg buffer and fixed on poly-lysine-coated slides using methanol at −20°C for 20 min, followed by acetone at −20°C for 10 min. The primary antibodies used were rabbit anti-PAR-2 (Hoege et al., 2010), rabbit anti-PAR-1 (Gonczy et al., 2001), rabbit anti-PKC-3 (Tabuse et al., 1998), mouse anti-PAR-3 (P4A1, DSHB), mouse anti-PGL-1 (K76, DSHB), and mouse anti-MEX-5 (Griffin et al., 2011; Schubert et al., 2000). Secondary antibodies used were goat anti-rabbit coupled to Alexa488, goat anti-rabbit coupled to Cy3, or goat anti-mouse coupled to Cy3 or Cy5, all at a 1:8,000 dilution. Samples were mounted with Vectashield Antifade Medium with DAPI (Vector Laboratories) to stain DNA. All antibodies used in this study are listed in Table S3.

### Quantification of cortical PAR proteins and cytoplasmic fate determinants

To quantify the distribution of PAR proteins at the cell cortex in live zygotes (Figure S2), we used ImaEdge software (Zhang et al., 2017), which was designed for automatic extraction of the cortex region from each frame of time-lapse movies. The cortical region along the entire circumference was divided into 100 sampling windows. The maximum intensity in each sampling window was used to represent the amount of proteins-of-interest at the cortex. We then generated a 2D heat map with the frame number as the horizontal axis and the position of the sampling windows as the vertical axis to integrate the spatiotemporal information of cortical PAR proteins.

The total levels of GFP::PAR-2 in zygotes were estimated by the average fluorescence intensities of GFP::PAR-2 in the cytoplasm of zygotes shortly after fertilization (before polarization of GFP::PAR-2 at the cortex). A box of 75.1 μm^2^ at the center position in the GFP::PAR-2 images (taken at about 2 μm below the upper lateral cortex) was used to measure the average intensity. The GFP::PAR-2 intensity of each zygote was normalized to the mean value of GFP::PAR-2 intensity in embryos expressing unmodified *gfp::par-2* transgene (JH2952).

The sizes of cortical GFP::PAR-2 and mCherry::PAR-6 domains were determined by the extent of each fluorescence protein occupying the circumference of the zygote. The surface of the zygote was manually traced using ImageJ software. The edge of each cortical domain was defined at the region where either GFP::PAR-2 or mCherry::PAR-6 intensity is about 40% of the maximum intensity within the corresponding cortical domain. The length of the cortical PAR-2 domain and that of the cortical PAR-6 domain were independently measured and represented as a percentage of the total circumference of the zygote.

The distribution of cortical PAR proteins (PAR-1, GFP::PAR-2, PAR-3, and mCherry::PAR-6) in zygotes were assessed by the integrated intensity of cortical PAR proteins within the anterior and the posterior cortical domains. The surface of the zygote was manually traced and divided into two halves with Metamorph software using the line-scan function. The segregation of the posterior proteins (PAR-1 and GFP::PAR-2) was represented as a ratio of their average intensity within the posterior domain to that within the anterior domain. The segregation of the anterior proteins (PAR-3 and mCherry::PAR-6) was represented as a ratio of their average intensity within the anterior domain to that within the posterior domain.

The distribution of cytoplasmic fate determinants, MEX-5 and PIE-1, were assessed by their average intensity within the anteromedial and the posteromedial cytoplasm, respectively. A box of 75.1 μm^2^ or 27.0 μm^2^ was used to measure the intensities of MEX-5 and PIE-1, respectively, with Metamorph software. The segregation of MEX-5 was represented as a ratio of the average intensity within the anteromedial cytoplasm over that within the posteromedial cytoplasm. The segregation of PIE-1 was represented as a ratio of the average intensity within the posteromedial cytoplasm over that within the anteromedial cytoplasm.

### Statistical tests and reproducibility

All statistical tests were performed using GraphPad Prism 7.0. All results presented in graphs represent the mean ± s.d. The D’Agostino-Pearson omnibus test was used for normality testing. A student’s t test (two-tailed distribution) and non-parametric Mann–Whitney U-test were used to calculate *p*-values. The exact sample number value is indicated in the corresponding figure or figure legend. All experiments with or without quantification were independently repeated at least three times with similar results, and the representative data are shown. No statistical method was used to predetermine the sample size. The experiments were not randomized. The investigators were not blinded to allocation during experiments or outcome assessment.

### Mathematical modeling of PAR polarity in C. elegans zygotes

#### 1) Previous models of PAR polarity in *C. elegans* zygotes

Previous models (Blanchoud et al., 2015; Dawes and Munro, 2011; Goehring et al., 2011) considered two species of PAR proteins. The anterior group of proteins (A) comprises of PAR-3, PAR-6, and PKC-3. The posterior group of proteins (P) is consisted of PAR-1, PAR-2, and LGL-1 (Figure S4A). The local cortical concentrations of A and P at time t and cortical position x, were designated by 𝐴 and 𝑃, respectively, and were calculated from the following equations:

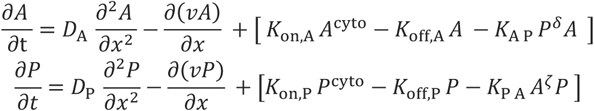

*D*_A_ and *D*_P_ represent the diffusivity of each PAR species in the membrane-bound state. *v*(*x,t*) refers to the cortical flow velocity. *k*_on,A_, *k*_off,A_, *k*_on,P_ and *k*_off,P_ are values for the respective coefficients for cortical association and dissociation of each PAR species. *A*^cyto^ and *P*^cyto^ are uniform cytoplasmic concentrations of A and P proteins, respectively. The second terms on the right side describe the initial segregation of both PAR species, which is mediated by advective cortical flow from the posterior pole to the anterior pole.

The above-described models (Blanchoud et al., 2015; Dawes and Munro, 2011; Goehring et al., 2011) reproduced the establishment and the maintenance of the steady-state PAR species distribution in wild-type zygotes. The model by (Goehring et al., 2011) also reproduced the PAR patterns in *spd-5(RNAi)* zygotes where advective cortical flows were attenuated. It also recapitulated the shift of a boundary position between two cortical domains in zygotes where either A or P was overexpressed or knocked-down (Goehring et al., 2011). However, these models were unable to reproduce the PAR distributions in zygotes wherein PAR-2 was predominant (by a combination of GFP::PAR-2 overexpression and *par-6(pRNAi)* treatment) (Figure S4C).

#### 2) A revised model of PAR polarity in *C. elegans* zygotes

To explain our observations and include findings from recent literatures, we revised the previous models by incorporating two additional PAR species, [CDC-42–PAR-6–PKC-3] and [CHIN-1], and including several molecular interactions among the PAR species (Figures 4A and S4B). PAR-3 recruits PAR-6 and PKC-3 to cortical cluster structures during polarization (Wang et al., 2017). CDC-42 can recruit PAR-6 and PKC-3 to the cortex independently of PAR-3 clusters (Rodriguez et al., 2017; Wang et al., 2017). Hence, the anterior cortical domain contains at least two species, [PAR-3–PAR-6–PKC-3] and [CDC-42– PAR-6–PKC-3]. CHIN-1 has been shown to localize at the posterior cortex and restricts activation of CDC-42 at the anterior cortical domain (Kumfer et al., 2010)(Sailer et al., 2015). PAR-2 is essential to recruit PAR-1 (Boyd et al., 1996) but dispensable to localize CHIN-1 at the posterior cortex (Sailer et al., 2015). CHIN-1 and PAR-1–PAR-2 act in parallel to maintain the polarized PAR domains (Sailer et al., 2015). Thus, the posterior cortical domain includes at least two species, [CHIN-1] and [PAR-1–PAR-2]. We did not include another posterior protein, LGL-1, because it is dispensable to pattern the cortical PAR domains (Beatty et al., 2010; Hoege et al., 2010). Hence, our revised model for the establishment of PAR polarity relies on four species of PAR proteins. [CDC-42–PAR-6–PKC-3] and [PAR-3–PAR-6– PKC-3] are referred as A_1_ and A_2_, respectively, as they localize at the anterior cortical domain. [CHIN-1] and [PAR-1–PAR-2] are referred as P_1_ and P_2_, respectively, as they localize at the posterior cortical domain.

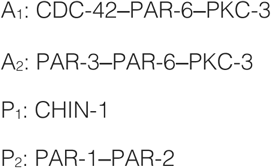

Given that CDC-42 is essential for PKC-3 to exclude PAR-2 independently of PAR-3 (Rodriguez et al., 2017), we included an inhibitory interaction from A_1_ [CDC-42–PAR-6–PKC-3] to P_2_ [PAR-1–PAR-2] in our model. Two previous reports (Rodriguez et al., 2017; Wang et al., 2017), also proposed that PAR-6 and PKC-3 could associate with either PAR-3 or CDC-42 on the cortex. Because PAR-3 is generally essential to recruit PAR-6 and PKC-3 during polarization (Wang et al., 2017), we include a positive interaction from A_2_ [PAR-3–PAR-6–PKC-3] to A_1_ [CDC-42–PAR-6–PKC-3]. CHIN-1 is excluded from the anterior cortex through an unknown mechanism that relies on PKC-3 (Sailer et al., 2015), and interferes active CDC-42 within the posterior cortical domain (Kumfer et al., 2010; Sailer et al., 2015). We thus include reciprocal inhibitory interactions: one from A_1_ [CDC-42–PAR-6–PKC-3] to P_1_ [CHIN-1], the other from P_1_ [CHIN-1] to A_1_ [CDC-42–PAR-6–PKC-3].

Diffusion rates of all four species in the cytoplasm are assumed to be much faster than their respective rates on the cell cortex (Goehring et al., 2011). Therefore, cytoplasmic concentration of these species should be uniform and quasi-statically adapted to their molecular interactions on the cortex. We denoted the cortical concentration of four species as *A*_1_,*A*_2_,*P*_1_,*P*_2_. The time evolution of the local concentration on the membrane at time *t* and position *x* is described by the following equations:

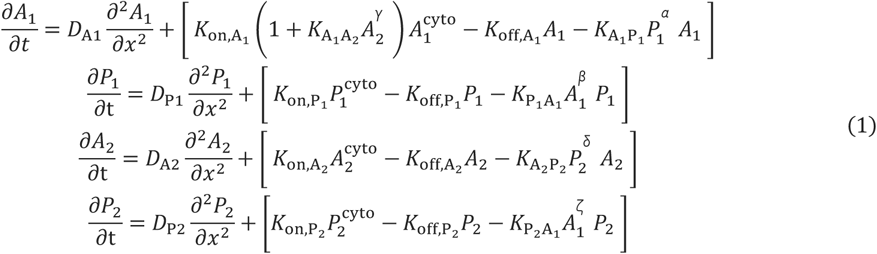

On the right side, the first terms are diffusion terms describing diffusional transport on the cortex with diffusion constants, *D*_A_1__, *D*_P_1__, *D*_A_2__ and *D*_P_2__. The next three terms are reaction terms explaining molecular interactions among the PAR species. The second set of terms describe cortical binding with binding rates *K*_on,A_1__,*K*_on,P_1__,*K*_on,A_2__,*K*_on,A_2__ and *K*_on,P_2__ per unit cytosolic concentrations. For example, in the first equation, the binding of A_1_ is promoted by the presence of A_2_ with the coefficient *K*_A_1_,A_2__. The third set of terms explain dissociation rates from the cortex with the unbinding rates *K*_off,A_1__,*K*_off,P_1__,*K*_off,A_2__ and *K*_off,A_2__. The fourth set of terms also refer to dissociation stimulated by the mutual antagonism between the molecules P_1_ and A_1_ with the rates *K*_A_1_P_1__ and *K*_A_1_P_1__, the inhibition from P_2_ to A_2_ with the rate *K*_A_2_P_2__, and the inhibition from A_1_ to P_2_ with the rate *K*_P_2_A_1__. Cytosolic concentrations of each PAR species were calculated from total concentration minus cortical concentration. For example, cytosolic concentrations of A_1_ can be described as following:

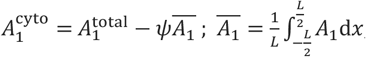

*Ψ* is the surface-to-volume conversion factor. *L* is the perimeter along the anteroposterior axis of zygote (Goehring et al., 2011). Since the periphery of the zygote was assumed to be a one-dimensional space, equations (1) was considered under the periodic boundary condition. We used Matlab to manually obtain numerical values of steady-state solutions in equations (1). For the initial condition, we considered a spatially uniform steady-state solution for the four variables and put higher values for P_1_ and P_2_ in the posterior region. Depending on the parameter sets, various PAR species (*A*_1_,*P*_1_,*A*_2_,*P*_2_) can distribute either uniformly or asymmetrically in a specific region of the cortex. The parameter values in wild-type (WT) zygotes are summarized in Table S4.

Like the previous models (Blanchoud et al., 2015; Dawes and Munro, 2011; Goehring et al., 2011), our model also requires multi-stability in the reaction terms that allows each PAR species to account for both the unpolarized and the polarized state at the cortex. Given that our modelling did not consider the advection term and relied on both the diffusion term and the reaction term, our simulation only constructs the steady-state distribution of each PAR species in zygotes.

The steady-state concentration profiles in WT condition are shown in Figure 4B. A_1_[CDC-42– PAR-6–PKC-3] and A_2_[PAR-3–PAR-6–PKC3] are enriched in the anterior domain, whereas the posterior domain shows high concentrations of P_1_[CHIN-1] and P_1_[PAR-1–PAR-2]. Hereafter, we normalized concentrations (*A*_1_,*A*_2_) and (*P*_1_,*P*_2_) with *A*_max_ and *P*_max_, respectively. *A*_max_ and *P*_max_ are the maximum values of cortical concentration in *A*_1_ and *A*_2_. and *P*_1_, and in *P*_2_, respectively. We then plotted normalized concentrations of the respective proteins against the length of the embryo, from the anterior to the posterior poles. The cortical domain size of each PAR species is defined as the percentage of the cortical region where protein concentration is equal to or greater than 40% of its maximum concentration at the cortex. The domain size of A_1_ species, for example, can be calculated by using the following formula:

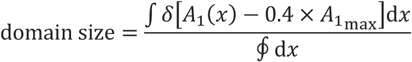

where the cortical concentration of A_1_ at the cortical position x is defined as *A*_1_(*x*), δ[*x*] is the Heaviside step function, and ∮ *dx* = *L*.

#### 3) Overexpression, knockdown, and knockout experiments *in silico*

To simulate experimental conditions where one (or more) PAR species was either knocked-down (KD), knocked-out (KO), or overexpressed (OE), we changed total concentrations of each PAR species in our simulations as they would have experimentally. The results for *in silico* KD, KO, OE manipulations are explained in detail below, and are shown in Figures 4C-4I and S4D-S4G.

##### 1. PAR-6 KO or PAR-2 KO

When the total concentration of A_1_ 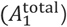 or P_2_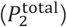 is at zero, cortical polarity exhibits nearly uniform. distribution of P_2_ or A_1_, respectively (Figures 4C and 4D). These results are consistent with previous observations (Boyd et al., 1996; Cuenca et al., 2003; Hung and Kemphues, 1999) and our results in Figure 2B.

##### 2. PAR-2 KD or PAR-2 OE

When 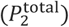 is reduced from the WT value of 1 to 0.1, the size of cortical P_2_ domain decreased, and that of cortical A_1_ domain increased, shifting the boundary between the A_1_ and the P_2_ domains from anterior to posterior (Figures 4F and S4D). When the total concentration of P_2_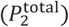 is increased from the WT value of 1 to 5, the size of cortical P_2_ domain increased, and that of cortical A_1_ domain reduced, shifting the boundary between the A_1_ and the P_2_ domains from posterior to anterior (Figures S4D and S4E). These results are consistent with the previous observation (Goehring et al., 2011) and our results in Figure 2A.

##### 3. PAR-2 OE either at WT background or at PAR-6 KD background

To systematically explore the effect of different levels of PAR-2 OE *in silico*, the total concentration of P_2_ 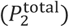 is increased from the WT value of 1 to 10. The corresponding size of A and P domains are shown in Figures S4D and S4E. The changes in cortical domain sizes in response to PAR-2 OE exhibit two distinct phases: When 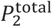 increases from 1, the size of A_1_ domain gradually decreases due to the expansion of P_2_ domain. At around 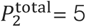 the solution of equations (1) shows a bifurcation, leading to a qualitative change in the distribution of concentrations. At this bifurcation point, distribution of P_2_ changes from the polarized state to the unpolarized state showing high concentration of P_2_ throughout the cortex. Despite the P_2_ distribution throughout the cortex, the distribution of A_1_ remains at the polarized state (Figures S4D and S4E).

We next tested if PAR-6 KD could compromise the stability of the polarized state of PAR-2 in response to PAR-2 OE treatment. When 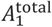 is lower than the WT value, the increase in 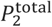 causes a similar bifurcation for P_2_ (the distribution changes from the polarized state to the unpolarized state) at a lower value of 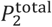 compared to the WT case (Figures 4H and S4F). Such a shift of the bifurcation point in 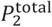 can be explained by the antagonistic interaction from A_1_ to P2. Because A_1_ antagonizes P_2_, a lower level of 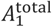 permits P_2_ to dominate the entire cortex even at lower 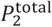. This result is consistent. with our observation of *par-6(pRNAi)* zygotes overexpressing GFP::PAR-2 in Figure 2A.

##### 4. CHIN-1 KD with or without PAR-2 OE

At the WT background, a reduction in 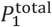 does not cause significant changes in the distribution patterns in A_1_, A_2_, and P_2_ (Figure 4E). In contrast, under P_2_ OE condition, reduction in 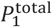 depolarizes A_1_ domain at the P_2_ bifurcation point where P_2_ becomes uniform at the cortex (Figures 4I and S4G). This result suggested a role of CHIN-1 in maintaining cortical PAR-6 domain in zygotes with a PAR-2 predominance. This is consistent with our observation of *chin-1(RNAi);par-6(pRNAi)* zygotes overexpressing GFP::PAR-2 in Figure 3F.

#### Mathematical modeling of PAR polarity in two-cell stage embryos

To simulate the distribution of PAR proteins in two-cell stage embryos, we applied the inter-connected network involving the four PAR species (A_1_, A_2_, P_1_, and P_2_) in Eq. (1), which was used for the PAR pattern simulation in zygotes, to AB and P1 cells. We considered simulation data of the steady-state distribution of the PAR species in zygote for the initial condition of two-cell stage embryos. The PAR species were compartmentalized between AB and P1 cells at the position of a cleavage furrow, which was experimentally measured in zygotes. Therefore, the initial profile of the PAR species in AB and P1 cells were copied respectively from the anterior and posterior portion of the zygote with length *L*_*AB*_ and *L*_*P*1_(*L*_*AB*_ + *L*_*P*1_ = *L*, the total length of the zygote). Based on *in vivo* observations, we set *L*_*AB*_ = 0.55 × *L* and *L*_*P*1_ = 0.45 × *L* for both wild-type and PAR-2-predominant embryos and *L*_*AB*_ = *L*_*P*1_ = 0.50 × *L* for *cye-1(RNAi)* embryos. The parameter values in wild-type (WT), P2OE, and *cye-1(RNAi)* embryos are summarized in Table S4, S5 and S6, respectively.

##### (1) Wild-type embryos

Through unequal inheritance of all four PAR species during the first cell division, AB and P1 cells inherited different total concentration of the PAR species. The AB cell increased the total concentration of A_1_ and A_2_ and decreased the total concentration of P_1_ and P_2_. The total concentration of the PAR species in the P_1_ cell were determined according to the law of mass conservation. Therefore, the P_1_ cell decreased the total concentration of A_1_ and A_2_ and increased the total concentration of P_1_ and P_2_. For the zygote and its daughter cells, the law of mass conservation was given as

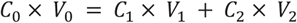

where C_0_, C_1_ and C_2_ are the total PAR concentration (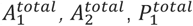 or 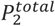) in zygote, AB cell,. and P_1_ cell, respectively. *V*_0_,*V*_1_, and *V*_2_ are their respective volumes. For the wild-type conditions, *V*_1_/*V*_0_ = 0.55 and *V*_2_/*V*_0_ 0.45. With the given changes in the total concentration of A_1_, A_2_, P_1_ and P_2_, the distribution of the PAR species in AB and P_1_ cells were numerically calculated until their spatial pattern reached the steady state. The parameter values used for WT embryo are given in Table S4.

Simulations in other conditions, such as PAR-2 overexpression (P2OE) and *cye-1(RNAi)* with higher or lower levels of PAR-2, were performed in a similar approach. The particular changes from simulations in the wild-type condition are summarized below.

##### (2) P2OE embryos

A higher total concentration of P_2_ was used for simulation of the steady state in zygotes. Because of nearly symmetric distribution of A_2_ and P_2_ in the P2OE zygotes, the total concentration of A_2_ and P_2_ in AB and P_1_ cells were similar to those in the zygote. The total concentration of A_1_ was increased and that of P_1_ was decreased for the AB cell, while the total concentration of A_1_ was decreased and that of P_1_ was increased for the P_1_ cell, satisfying the conservation law. The parameter values are summarized in Table S5.

##### (3) *cye-1(RNAi)* embryos

Because of equal inheritance of all PAR proteins between the *cye-1(RNAi)* daughter cells, the total concentrations of A_1_, A_2_, P_1_, and P_2_ in these cells were the same as those in the zygote. For the initial condition of the daughter cells, we considered a spatially uniform steady state solution for the four variables (*A*_1_, *A*_2_, *P*_1_, and *P*_2_) with random fluctuations. The total concentration of P_2_ was modified to simulate the cells with higher or lower P_2_ condition. The parameter values are summarized in Table S6.

### Data availability

All data and materials supporting the findings of this study are available from the corresponding author on reasonable request.

