## Supplementary figures and tables for "Binary decision between asymmetric and symmetric cell division is defined by the balance of PAR proteins in *C. elegans* embryos"

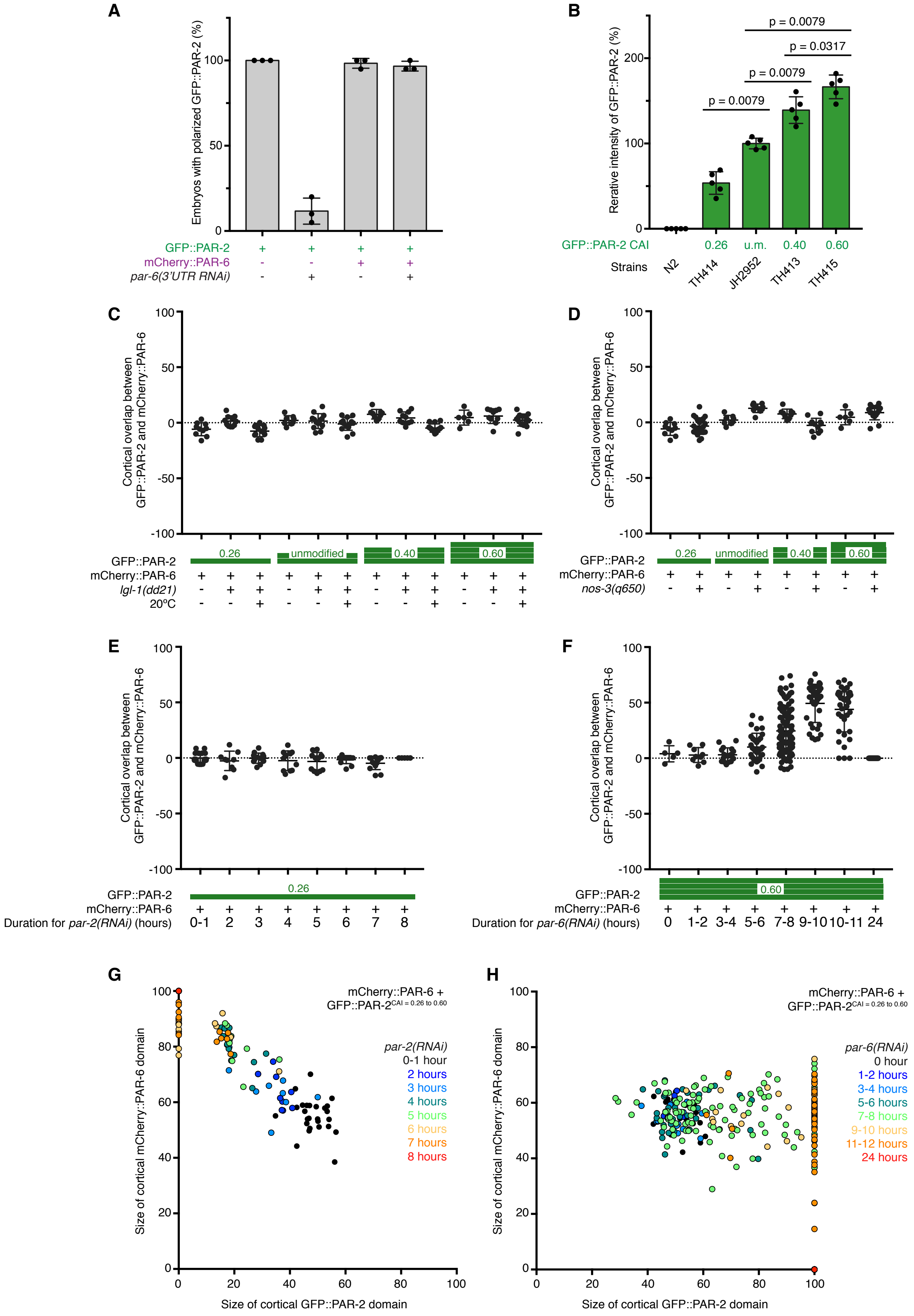


### Figure S1

Binary decision between asymmetric and symmetric cell division is defined by the balance of PAR proteins in C. elegans embryos

Yen Wei Lim, Fu-Lai Wen, Prabhat Shankar, Tatsuo Shibata, Fumio Motegi

**Figure S1. The landscape of PAR polarity patterning in *C. elegans* zygotes.**

(A) The graph indicates the percentage of zygotes that polarized GFP::PAR-2 at the posterior cortex. Control zygotes and zygotes expressing mCherry::PAR-6 were imaged with or without *par-6(3'-UTR RNAi)* treatment. Each experiment was repeated three times and analyzed 20 zygotes.

(B) The graph shows the relative fluorescence intensities of cytoplasmic GFP::PAR-2 in zygotes before polarization. The GFP::PAR-2 intensity in each zygote was normalized to the mean value of GFP::PAR-2 intensity in embryos expressing unmodified *gfp::par-2* transgene. Data presented as mean ± s.d. from n = 5 zygotes. *p*-values; Mann–Whitney test.

(C and D) The graphs depict the size of cortical domains where mCherry::PAR-6 and GFP::PAR-2 overlapped. (C) Data presented as mean ± s.d. from n = 10, 11, 13, 8, 13, 14, 9, 10, 14, 6, 10, 15 zygotes.

(D) Data presented as mean ± s.d. from n = 10, 22, 8, 10, 9, 12, 6, 15 zygotes.

(E and F) Exploration of the limit of PAR proteins segregation by time-course analysis of *par-2(pRNAi)* zygotes (E) and *par-6(pRNAi)* zygotes (F). The graphs depict the size of cortical GFP::PAR-2 domain before and after RNAi treatment. (E) Data presented as mean ± s.d. from n = 9, 9, 13, 11, 12, 13, 13, 5 zygotes. (F) Data presented as mean ± s.d. from n = 5, 9, 18, 33, 105, 36, 38, 21 zygotes.

(C-F) Height of green bars indicate the *gfp::par-2* transgene CAI.

(G and H) The landscape of PAR polarity patterning *in vivo*. The sizes of mCherry::PAR-6 and GFP::PAR-2 cortical domains in zygotes with various PAR protein balances are plotted. (G) Gradual depletion of PAR-2 by time-course analysis of *par-2(pRNAi)* zygotes caused a reduction of the GFP::PAR-2 domain size, followed by abrupt disappearance of the GFP::PAR-2 domain. (H) Gradual depletion of PAR-6 by time-course analysis of *par-6(pRNAi)* zygotes led to co-existence of the polarized mCherry::PAR-6 domain and the uniform GFP::PAR-2 domain at the cortex.

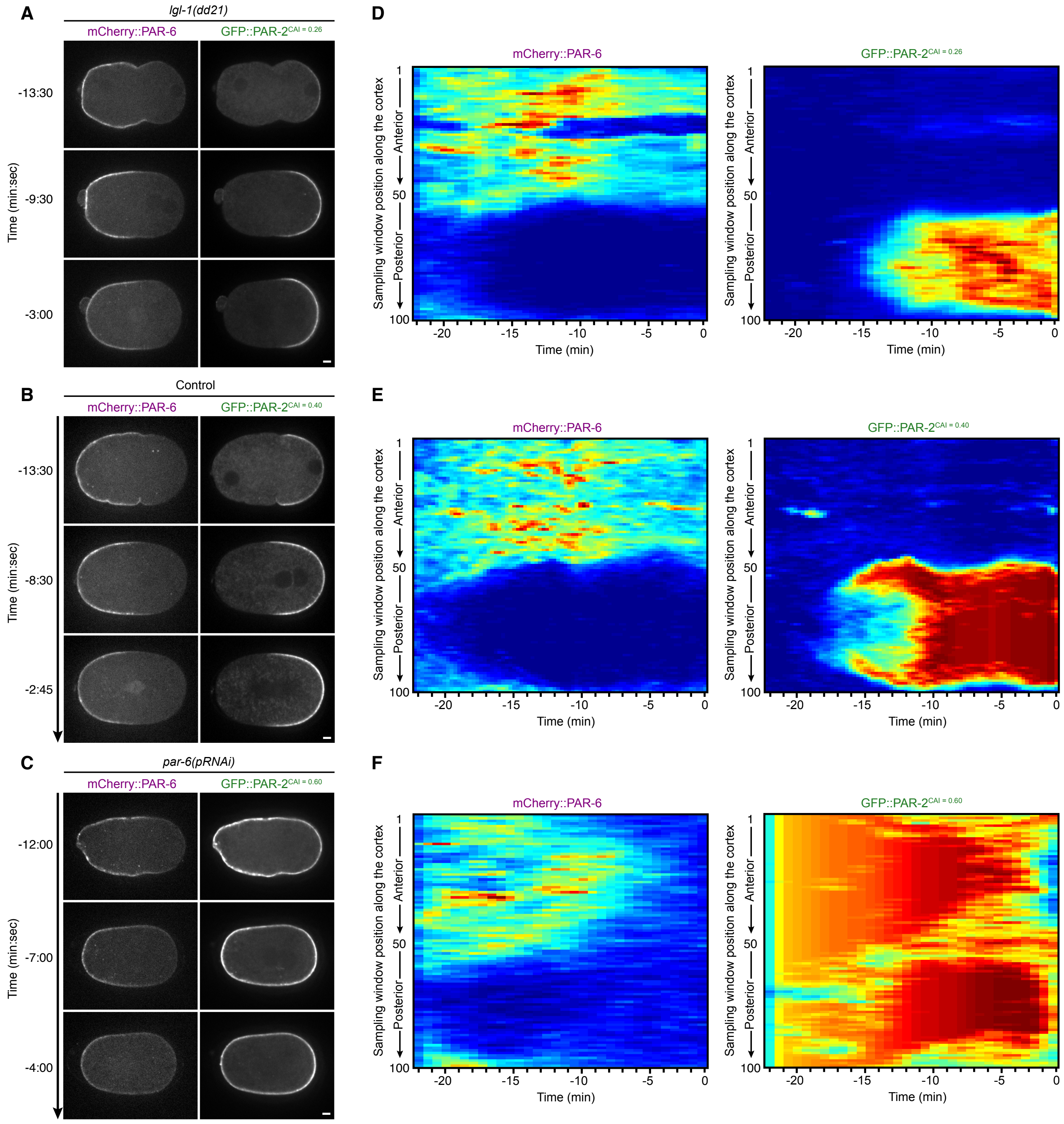


### Figure S2

Binary decision between asymmetric and symmetric cell division is defined by the balance of PAR proteins in C. elegans embryos

Yen Wei Lim, Fu-Lai Wen, Prabhat Shankar, Tatsuo Shibata, Fumio Motegi

**Figure S2. Dynamics of cortical PAR proteins in zygotes when the PAR balance is manipulated.**

(A-C) Representative time-lapse images of GFP::PAR-2 and mCherry::PAR-6 during polarization in a *lgl-1(dd21);par-2(ok1723)* zygote expressing GFP::PAR-2$^{CAI = 0.26}$ (A), a *par-2(ok1723)* zygote expressing GFP::PAR-2$^{CAI = 0.40}$ (B), and a *par-6(pRNAi);par-2(ok1723)* zygote expressing GFP::PAR-2$^{CAI = 0.60}$ (C). Scale bar, 5 μm.

(D-F) Representative kymographs of cortical intensity dynamics of GFP::PAR-2 and mCherry::PAR-6 in zygotes described in (A-C). The cell boundary position and the thickness of the cell cortex were defined by intensity-based segmentation with ImaEdge software. The segmented cortex was then divided into 100 equally-spaced side-by-side boxes. The maximum intensities of GFP::PAR-2 and mCherry::PAR-6 within each sampling box were visualized as a 2D heat map with the horizontal axis as the time frame and the vertical axis as the position of sampling boxes.

(A-F) The times stated are with respect to the onset of cytokinesis.

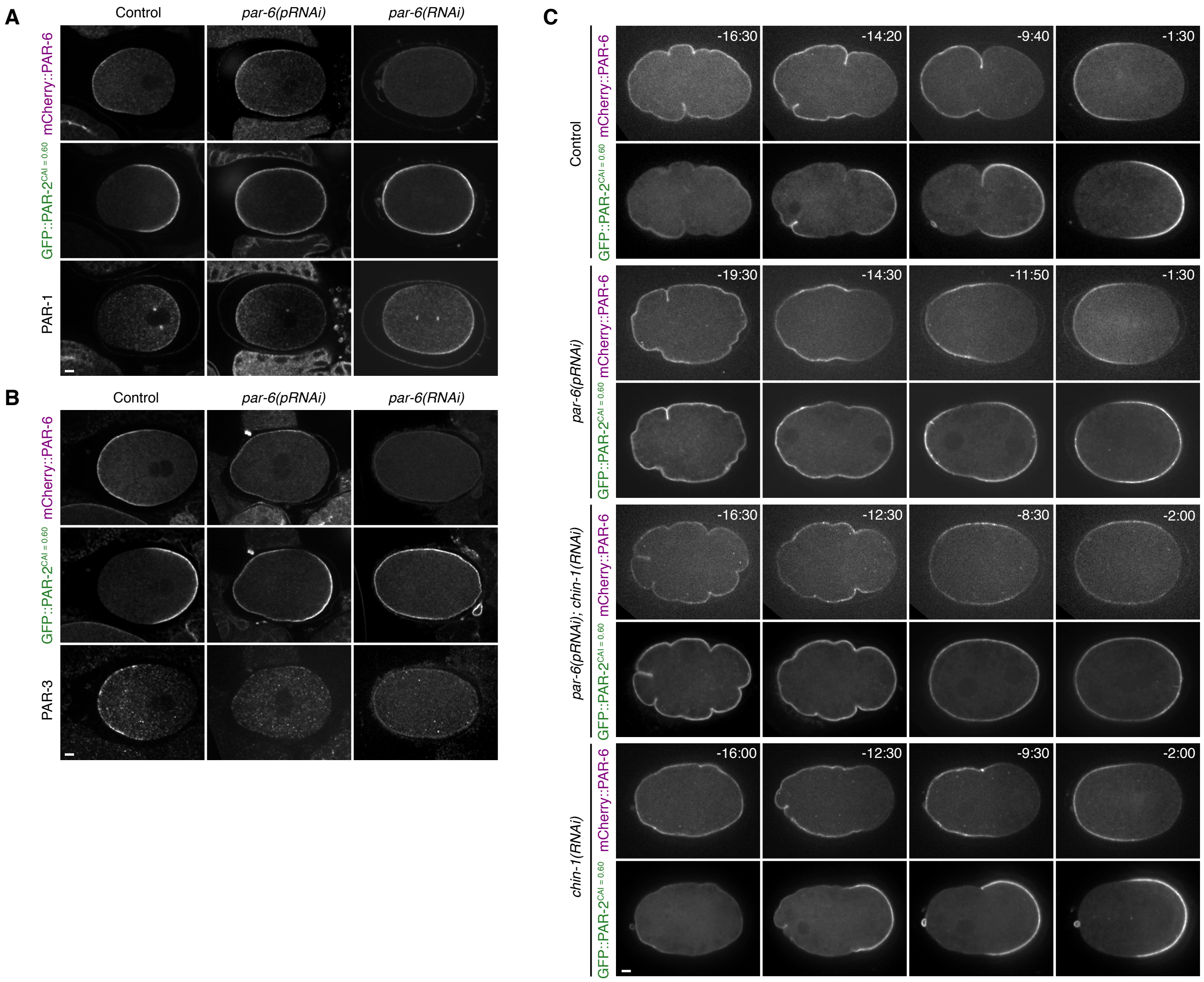


**Figure S3**

Binary decision between asymmetric and symmetric cell division is defined by the balance of PAR proteins in C. elegans embryos

Yen Wei Lim, Fu-Lai Wen, Prabhat Shankar, Tatsuo Shibata, Fumio Motegi

**Figure S3. A combinatorial network of two reciprocal exclusion pathways ensures cortical patterning of PAR proteins.**

(A) Representative images of mCherry::PAR-6, GFP::PAR-2, and PAR-1 in a control zygote and in *par-6(pRNAi)* zygotes wherein PAR-2 was predominant.

(B) Representative images of mCherry::PAR-6, GFP::PAR-2, and PAR-3 in a control zygote and in *par-6(pRNAi)* zygotes wherein PAR-2 was predominant.

(C) Representative time-lapse movies of zygotes under control, *par-6(pRNAi), par-6(pRNAi);chin-1(RNAi),* and *chin-1(RNAi)* conditions. The times stated (min:sec) are with respect to the onset of cytokinesis.

(A-C) Scale Bar, 5 μm.

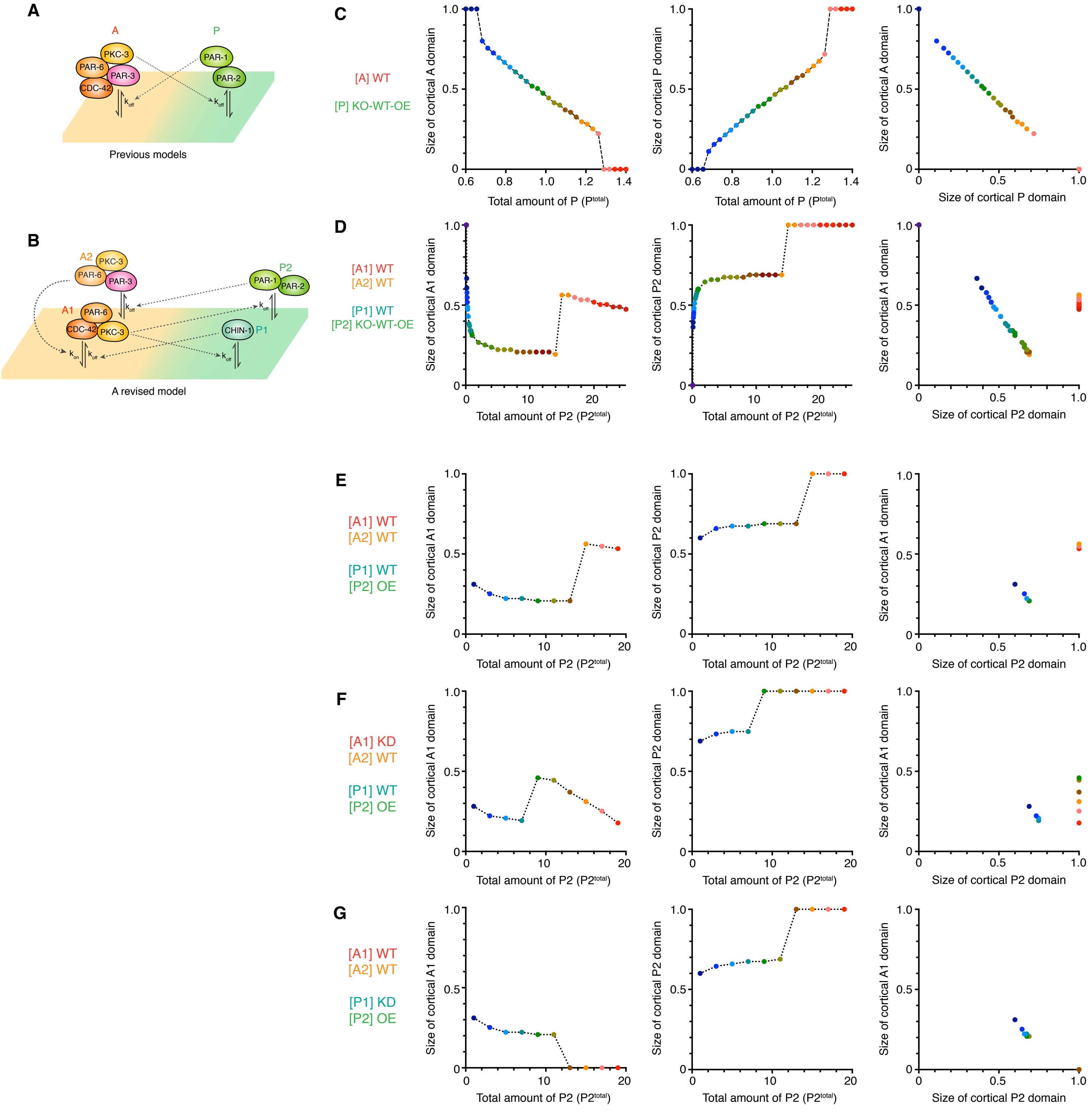


### Figure S4

Binary decision between asymmetric and symmetric cell division is defined by the balance of PAR proteins in C. elegans embryos

Yen Wei Lim, Fu-Lai Wen, Prabhat Shankar, Tatsuo Shibata, Fumio Motegi

Figure S4. The landscape of PAR polarity patterning *in silico.*

(A and B) Two models of PAR polarity patterning. (A) The conventional PAR network comprises a single mutual inhibition pathway between two PAR species, A [PAR-3–PAR-6–PKC-3–CDC-42] and P [PAR-1–PAR-2]. (B) The revised model consists of four PAR species, $A_1$ [CDC-42–PAR-6–PKC-3], $A_2$ [PAR-3–PAR-6–PKC-3], $P_1$ [CHIN-1], and $P_2$ [PAR-1–PAR-2].

(C) The sizes of A (left) and P (middle) cortical domains are plotted as a function of the total amount of P ($P^{total}$). A scatter plot of the sizes of these cortical domains is also shown (right).

(D-G) The sizes of A1 (left) and P2 (middle) cortical domains (d-f) are plotted as a function of the total amount of P2 ($P_2^{total}$). A scatter plot of the sizes of these cortical domains is also shown (right).

(D) When $P_2^{total}$ is reduced from the wild-type (WT) value 1 to 0.1, the size of cortical $P_2$ domain decreased, and that of cortical $A_1$ domain increased, shifting the $A_1$-$P_2$ domain boundary toward the posterior pole.

(E) When $P_2^{total}$ is increased from 1 to 5, the size of cortical $P_2$ domain increased, and that of cortical $A_1$ domain reduced, shifting the $A_1$-$P_2$ domain boundary toward the anterior pole. At around $P_2^{total}$=5, distribution of $P_2$ changes from the polarized state to the unpolarized state, showing uniform $P_2$ concentration throughout the cortex, while the distribution of $A_1$ remains at the polarized state.

(F) When the total amount of $A_1$ ($A_1^{total}$) is lower (62% of the WT value), the increase in $P_2^{total}$ causes a similar polarized-to-unpolarized shift for $P_2$ distribution at a lower value of $P_2^{total}$ compared to the WT case.

(G) Reduction in the total amount of $P_1$ ($P_1^{total}$) does not cause significant changes in the patterning of PAR species at the WT background, but it depolarizes the $A_1$ domain at the point where $P_2$ becomes uniform at the cortex.

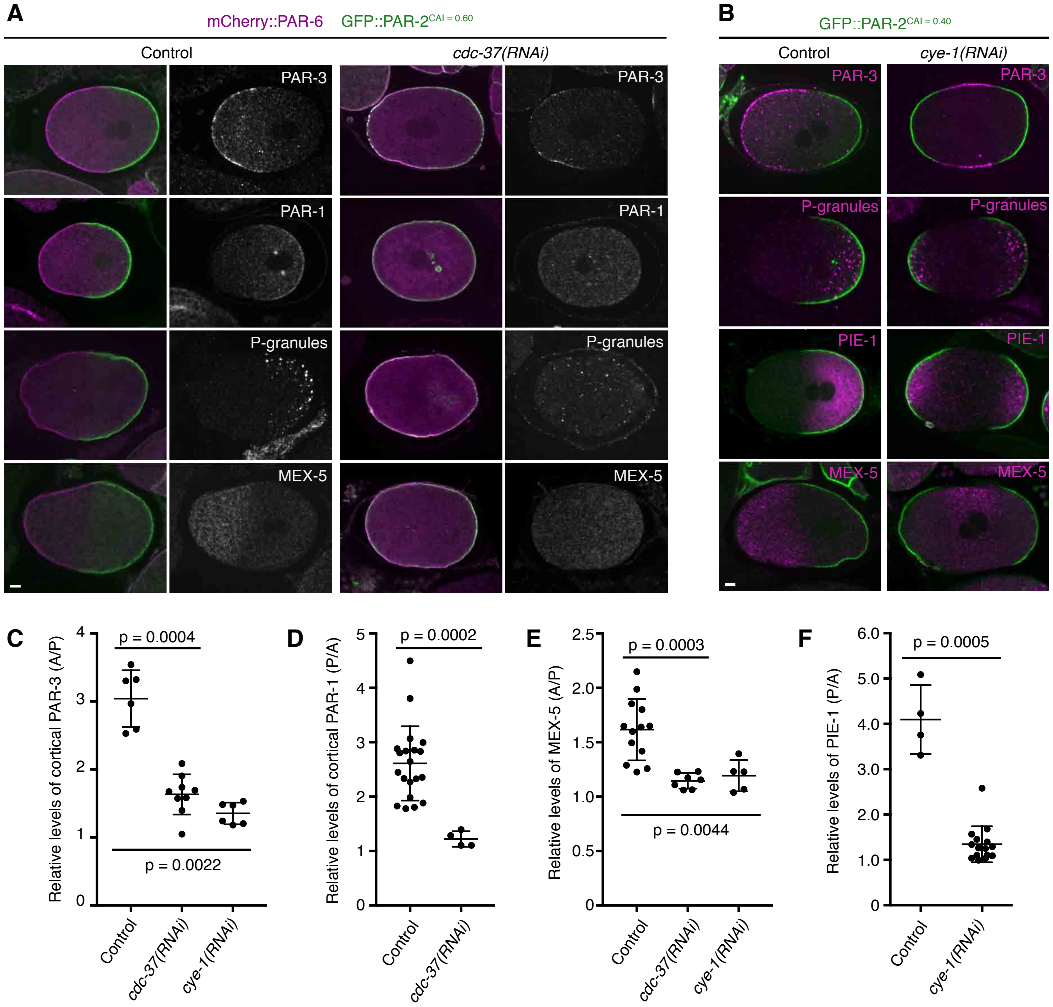


#### Figure S5

Binary decision between asymmetric and symmetric cell division is defined by the balance of PAR proteins in C. elegans embryos

Yen Wei Lim, Fu-Lai Wen, Prabhat Shankar, Tatsuo Shibata, Fumio Motegi

**Figure S5. Distribution of fate determinants in *cdc-37(RNAi)* and *cye-1(RNAi)* zygotes.**

(A) Representative images of the distribution of GFP::PAR-2$^{CAI = 0.60}$ (green), mCherry::PAR-6 (magenta), PAR-3, PAR-1, P-granules, and MEX-5 in zygotes under control and *cdc-37(RNAi)* conditions. Scale Bar, 5 μm.

(B) Representative images of the distribution of GFP::PAR-2$^{CAI = 0.40}$ (green), PAR-3, P-granules, PIE-1, and MEX-5 (magenta) in zygotes under control and *cye-1(RNAi)* conditions. Depletion of CYE-1 caused failures in unequal inheritance of PAR proteins (GFP::PAR-2 and PAR-3) and their effectors (P-granules, PIE-1, and MEX-5). Scale Bar, 5 μm.

(C) The graph depicts the relative levels of PAR-3 at the anterior cortex to the posterior cortex in zygotes under control, *cdc-37(RNAi),* and *cye-1(RNAi)* conditions. Data present mean ± s.d. from n = 6, 9, 6 zygotes.

(D) The graph depicts the relative levels of PAR-1 at the posterior cortex to the anterior cortex in zygotes under control and *cdc-37(RNAi)* conditions. Data present mean ± s.d. from n = 20, 4 zygotes.

(E) The graph depicts the relative levels of MEX-5 withih the anterior cytoplasm to the posterior cytoplasm in zygotes under control, *cdc-37(RNAi),* and *cye-1(RNAi)* conditions. Data present mean ± s.d. from n = 13, 7, 5 zygotes.

(F) The graph depicts the relative levels of PIE-1 within the posterior cytoplasm to the anterior cytoplasm in zygotes under control and *cye-1(RNAi)* conditions. Data present mean ± s.d. from n = 4, 15 zygotes.

(C-F) *p*-values; Mann–Whitney test.

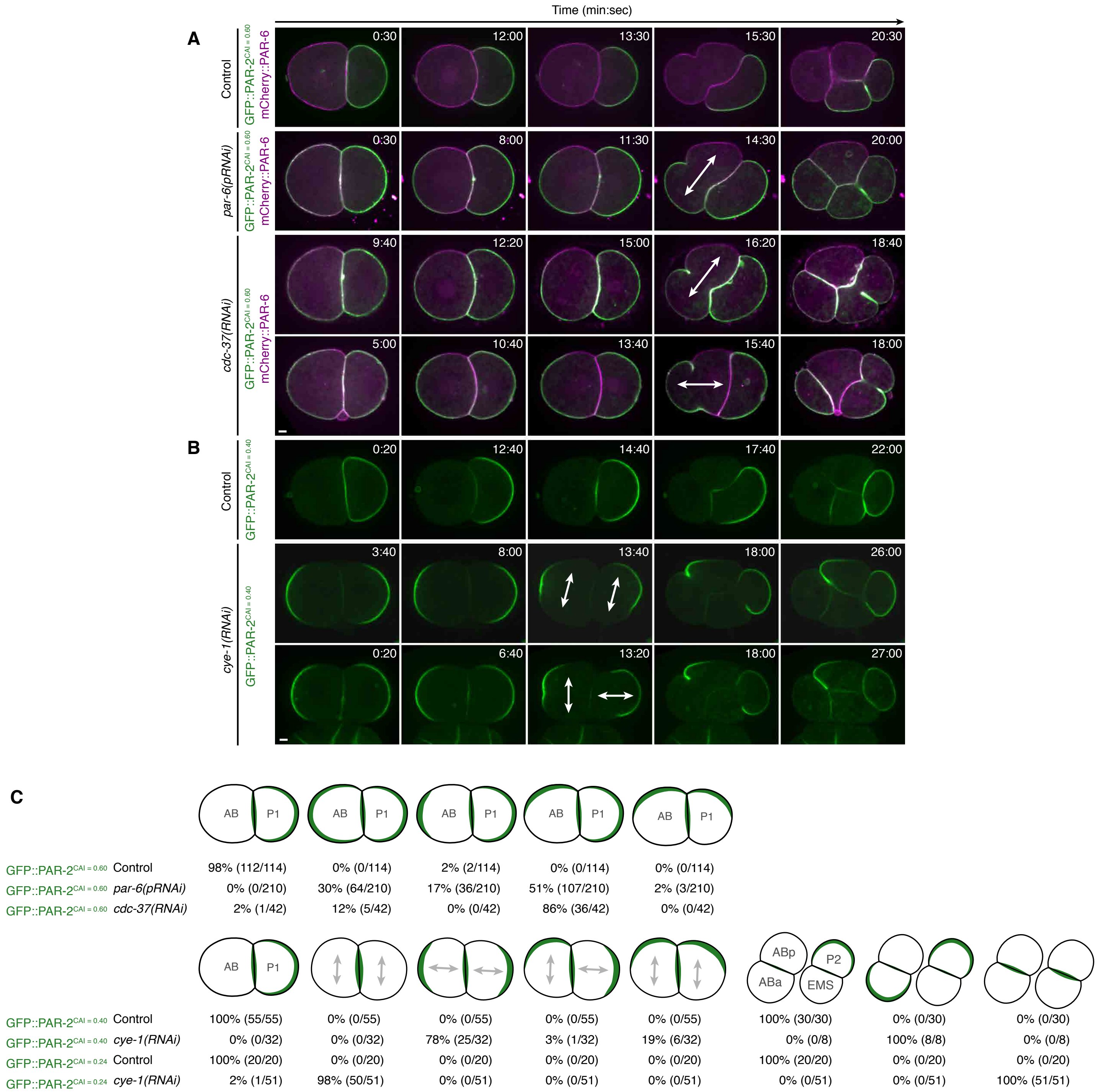


### Figure S6

Binary decision between asymmetric and symmetric cell division is defined by the balance of PAR proteins in C. elegans embryos

Yen Wei Lim, Fu-Lai Wen, Prabhat Shankar, Tatsuo Shibata, Fumio Motegi

**Figure S6. PAR polarity patterning in two-cell embryos in which the PAR balance is manipulated.**

(A) Representative time-lapse images of embryos expressing mCherry::PAR-6 (magenta) and either GFP::PAR-2$^{CAI = 0.40}$ or GFP::PAR-2$^{CAI = 0.60}$ (green) under control, *par-6(pRNAi), par-6(RNAi),* and *cdc-37(RNAi)* conditions. The PAR-2-predominant AB cells started to polarize both GFP::PAR-2 and mCherry::PAR-6 during mitosis, leading to the unequal inheritance of cortical GFP::PAR-2 and mCherry::PAR-6 between their daughter cells.

(B) Representative time-lapse images of embryos expressing GFP::PAR-2$^{CAI = 0.40}$ (green) under control and *cye-1(RNAi)* conditions. Both cells in two-cell stage *cye-1(RNAi)* embryos segregated GFP::PAR-2, resulting in the unequal inheritance of GFP::PAR-2 between their daughter cells.

(A and B) The times stated are with respect to the completion of the first cytokinesis. Scale Bar, 5 μm.

(C) The patterns of GFP::PAR-2 distribution in two-cell stage and four-cell stage embryos from the various conditions.

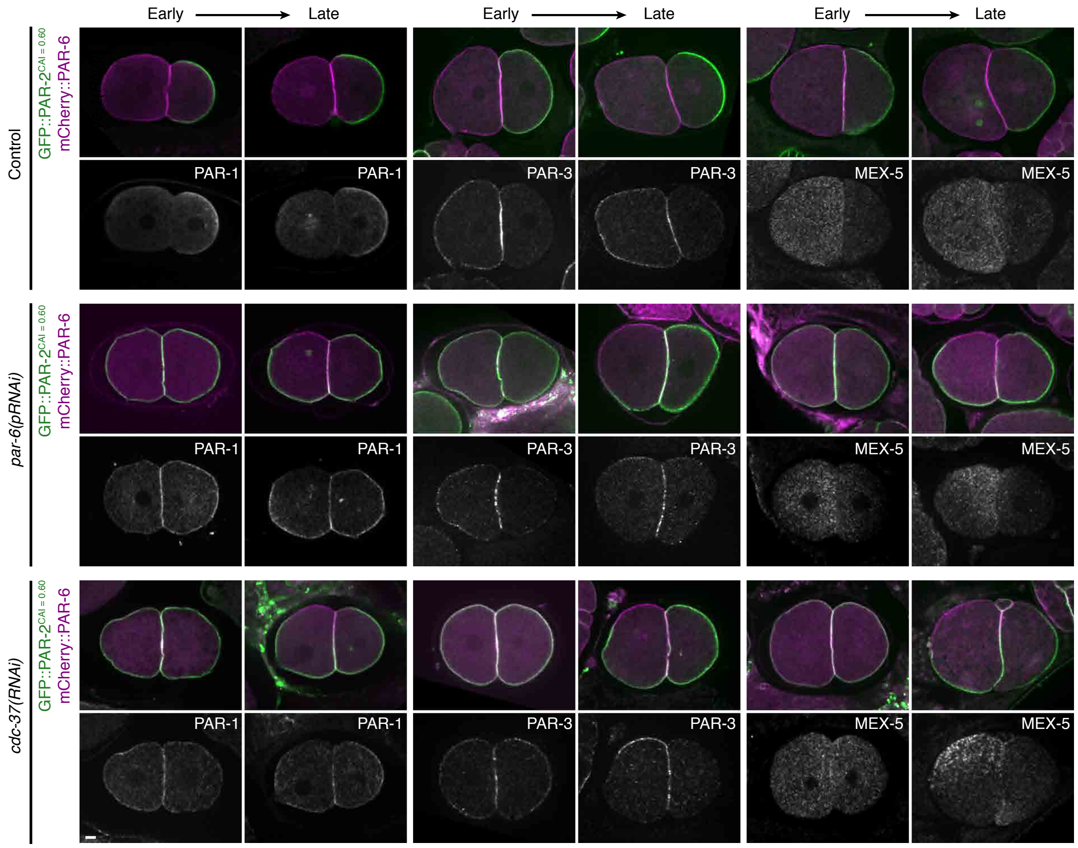


### Figure S7

Binary decision between asymmetric and symmetric cell division is defined by the balance of PAR proteins in C. elegans embryos

Yen Wei Lim, Fu-Lai Wen, Prabhat Shankar, Tatsuo Shibata, Fumio Motegi

**Figure S7. Patterns of fate determinants in 2-cell embryos in which the PAR balance is manipulated.**

Representative images of the distributions of PAR-1, PAR-3, and MEX-5 in embryos expressing both mCherry::PAR-6 (magenta) and GFP::PAR-2$^{CAI = 0.60}$ (green) under control, *par-6(pRNAi),* and *cdc-37(RNAi)* conditions. AB cells during early and late stages of mitosis are shown. Both cortical PAR proteins and cytoplasmic MEX-5 started to polarize during mitosis in PAR-2-predominant AB cells. Scale Bar, 5 μm.

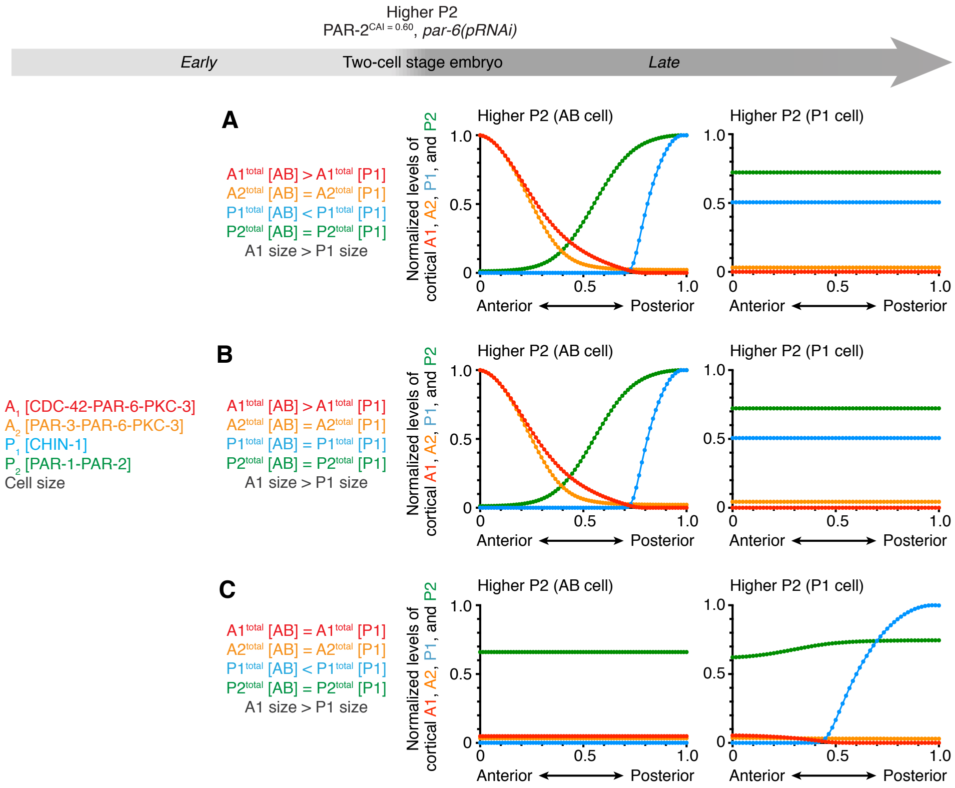


**Figure S8**

Binary decision between asymmetric and symmetric cell division is defined by the balance of PAR proteins in C. elegans embryos
Yen Wei Lim, Fu-Lai Wen, Prabhat Shankar, Tatsuo Shibata, Fumio Motegi

**Figure S8. Unequal inheritance of $A_1$ is essential for the PAR patterning in PAR-2-predominant embryos**

Steady-state analysis of the PAR network model in PAR-2-predominant embryos with or without unequal inheritance of $A_1$ or $P_1$ between AB and P1 cells. Predicted distributions of the PAR species at the cortex along the anteroposterior axis in an AB cell (left) and a P1 cell (right) with following conditions are shown.

(A) Unequal inheritance of $A_1$ and $P_1$ between AB and P1 cells.

($A_1^{total}[AB] > A_1^{total}[P1]$; $P_1^{total}[AB] < P_1^{total}[P1]$).

(B) Equal inheritance of $P_1$ between AB and P1 cells.

($A_1^{total}[AB] > A_1^{total}[P1]$; $P_1^{total}[AB] = P_1^{total}[P1]$).

(C) Equal inheritance of $A_1$ between AB and P1 cells.

($A_1^{total}[AB] = A_1^{total}[P1]$; $P_1^{total}[AB] < P_1^{total}[P1]$).

Table S1. ***C. elegans*** strains used in this study

| Strain | Genotype | Source |
|---|---|---|
| N2 | wild type | (Brenner, 1974) |
| JH2952 | *unc-119(ed3) III; temIs7[pIC26::par-2 re-coded], par-2(ok1723)* | (Motegi et al., 2011) |
| TH413 | *unc-119(ed3) III; ddIs25[gfp::par-2 re-coded, CAI=0.41]; par-2(ok1723)* | (Goehring et al., 2011) |
| TH414 | *unc-119(ed3) III; ddIs238[gfp::par-2 re-coded, CAI=0.26]; par-2(ok1723)* | (Goehring et al., 2011) |
| TH415 | *unc-119(ed3) III; ddIs239[gfp::par-2 re-coded, CAI=0.60]; par-2(ok1723)* | (Goehring et al., 2011) |
| MOT119 | *unc-119(ed3) III; ddIs238[gfp::par-2 re-coded, CAI=0.26]; par-2(ok1723); temIs17[pie-1p::mcherry::par-6::pie-1 3'UTR]* | (Zhang et al., 2017) |
| MOT121 | *unc-119(ed3) III; temIs7[pIC26::par-2 re-coded], par-2(ok1723); temIs17[pie-1p::mcherry::par-6::pie-1 3'UTR]* | (Zhang et al., 2017) |
| MOT118 | *unc-119(ed3) III; ddIs25[gfp::par-2 re-coded, CAI=0.41]; par-2(ok1723); temIs17[pie-1p::mcherry::par-6::pie-1 3'UTR]* | (Zhang et al., 2017) |
| MOT120 | *unc-119(ed3) III; ddIs239[gfp::par-2 re-coded, CAI=0.60]; par-2(ok1723); temIs17[pie-1p::mcherry::par-6::pie-1 3'UTR]* | (Zhang et al., 2017) |
| MOT401 | *unc-119(ed3) III; ddIs239[gfp::par-2 re-coded, CAI=0.60]; par-2(ok1723); ; temIs17[pie-1p::mcherry::par-6::pie-1 3'UTR]; axIs1464[pie-1p::gfp::pgl-3::pie-1 3'UTR]* | This study |
| MOT175 | *unc-119(ed3) III; temIs7[pIC26::par-2 re-coded], par-2(ok1723); nos-3(q650) II* | This study |
| MOT177 | *unc-119(ed3) III; ddIs25[gfp::par-2 re-coded, CAI=0.41]; par-2(ok1723); nos-3(q650) II* | This study |
| MOT179 | *unc-119(ed3) III; ddIs238[gfp::par-2 re-coded, CAI=0.26]; par-2(ok1723); nos-3(q650) II* | This study |
| MOT183 | *unc-119(ed3) III; ddIs239[gfp::par-2 re-coded, CAI=0.60]; par-2(ok1723); nos-3(q650) II* | This study |

| | | |
|---|---|---|
| MOT176 | *unc-119(ed3) III; ddIs238[gfp::par-2 re-coded, CAI=0.26]; par-2(ok1723); temIs17[pie-1p::mcherry::par-6::pie-1 3'UTR]; nos-3(q650) II* | This study |
| MOT178 | *unc-119(ed3) III; temIs7[pIC26::par-2 re-coded], par-2(ok1723); temIs17[pie-1p::mcherry::par-6::pie-1 3'UTR]; nos-3(q650) II* | This study |
| MOT180 | *unc-119(ed3) III; ddIs25[gfp::par-2 re-coded, CAI=0.41]; par-2(ok1723); temIs17[pie-1p::mcherry::par-6::pie-1 3'UTR]; nos-3(q650) II* | This study |
| MOT184 | *unc-119(ed3) III; ddIs239[gfp::par-2 re-coded, CAI=0.60]; par-2(ok1723); temIs17[pie-1p::mcherry::par-6::pie-1 3'UTR]; nos-3(q650) II* | This study |
| MOT192 | *unc-119(ed3) III; temIs7[pIC26::par-2 re-coded], par-2(ok1723); lgl-1(dd21) X* | This study |
| MOT190 | *unc-119(ed3) III; ddIs25[gfp::par-2 re-coded, CAI=0.41]; par-2(ok1723); lgl-1(dd21) X* | This study |
| MOT185 | *unc-119(ed3) III; ddIs238[gfp::par-2 re-coded, CAI=0.26]; par-2(ok1723); lgl-1(dd21) X* | This study |
| MOT199 | *unc-119(ed3) III; ddIs239[gfp::par-2 re-coded, CAI=0.60]; par-2(ok1723); lgl-1(dd21) X* | This study |
| MOT186 | *unc-119(ed3) III; ddIs238[gfp::par-2 re-coded, CAI=0.26]; par-2(ok1723); temIs17[pie-1p::mcherry::par-6::pie-1 3'UTR]; lgl-1(dd21) X* | This study |
| MOT188 | *unc-119(ed3) III; temIs7[pIC26::par-2 re-coded], par-2(ok1723); temIs17[pie-1p::mcherry::par-6::pie-1 3'UTR]; lgl-1(dd21) X* | This study |
| MOT189 | *unc-119(ed3) III; ddIs25[gfp::par-2 re-coded, CAI=0.41]; par-2(ok1723); temIs17[pie-1p::mcherry::par-6::pie-1 3'UTR]; lgl-1(dd21) X* | This study |
| MOT19Y | *unc-119(ed3) III; ddIs239[gfp::par-2 re-coded, CAI=0.60]; par-2(ok1723); temIs17[pie-1p::mcherry::par-6::pie-1 3'UTR]; lgl-1(dd21) X* | This study |
| JK2589 | *nos-3(q650) II* | (Pacquelet et al., 2008) |
| TH131 | *lgl-1(dd21) X* | (Hoege et al., 2010) |
| YAA16 | *unc-119(ed3) III; ddIs239[gfp::par-2 re-coded, CAI=0.60]; par-2(ok1723); temIs17[pie-1p::mcherry::par-6::pie-1 3'UTR]* | (Arata et al., 2016) |

Table S2. Materials for feeding RNAi experiments

| Target genes | Sequence | Source |
|---|---|---|
| ***par-1*** | H39E23.1 | SourceBioscience |
| ***par-2*** | F58B6.3 | (Motegi et al., 2011) |
| ***par-6*** | T26E3.3 | SourceBioscience |
| ***nos-3*** | Y53C12B.3 | SourceBioscience |
| ***lgl-1*** | F56F10.4 | SourceBioscience |
| ***chin-1*** | BE0003N10.2 | SourceBioscience |
| ***cdc-42*** | R07G3.1 | SourceBioscience |

Table S3. Antibodies used in this study

| Target protein | Sequence | Animals used for immunization | Notes | Source |
|---|---|---|---|---|
| PAR-1 | H39E23.1 | Rabbit | Antigen: His::PAR-1. 1:100 dilution was used. | (Hoege et al., 2010) |
| PAR-1 | H39E23.1 | Rabbit | Antigen: GST-tagged 780-1232 aa of PAR-1. 1:10,000 dilution was used. | (Gonczy et al., 2001) |
| PAR-2 | F58B6.3 | Rabbit | Antigen: His::PAR-2. 1:2,000 dilution was used. | (Hoege et al., 2010) |
| PAR-3 | F54E7.3 | Mouse | P4A1 from DHSB. 1/30 dilution was used. | DHSB |
| PAR-6 | T26E3.3 | Rabbit | 1:6,000 dilution was used. | (Labbe et al., 2006) |
| PAR-6 | T26E3.3 | Rabbit | Antigen: GST::PAR-6. Antibody was purified by NC blot-affinity purification. | This study |
| PKC-3 | F09E5.1 | Rabbit | 1:6,000 dilution was used. | (Sugiyama et al., 2008) |
| PGL-1 | ZK381.4 | Mouse | OIC1D4 from DHSB. 1:10 dilution was used. | DHSB |
| MEX-5 | W02A2.7 | Mouse | 1:500 dilution with 5% BSA was used. | (Schubert et al., 2000) |
| CHIN-1 | BE0003N10.2 | Rabbit | Antigen: 137-236 aa of CHIN-1. | (Kumfer et al., 2010) |
| CHIN-1 | BE0003N10.2 | Mouse | Antigen: a combination of 3 synthetic peptides. | Abmart: http://www.ab-mart.com/worm/anti-chin-1%20(Q965N4).html |
| Rabbit IgG | | Donkey | Cy5-labeled. 1:400 dilution was used. | Jackson (711-175-152) |
| Mouse IgG | | Donkey | Cy5-labeled. 1:400 dilution was used. | Jackson (711-175-150) |

| | | | |
|---|---|---|---|
| Guinea pig IgG | Donkey | Cy5-labeled. 1:400 dilution was used. | Jackson (706-175-148) |
| Rabbit IgG | Goat | Cy3-labeled. 1:8,000 dilution was used. | Abcam (ab6939) |
| Mouse IgG | Goat | Cy3-labeled. 1:8,000 dilution was used. | Abcam (ab97035) |
| Rabbit IgG | Goat | Alexa Fluor 594-labeled. 1:8,000 dilution was used. | Lab-Chem Enzyme (A11005) |
| Mouse IgG | Goat | Alexa Fluor 594-labeled. 1:8,000 dilution was used. | Lab-Chem Enzyme (A11012) |

2

Table S4. Parameter values used in the simulations of cortical PAR polarity in *C. elegans* embryos

| Parameter | Value | Source |
|---|---|---|
| **$A_1$ (CDC-42–PAR-6–PKC-3)** | | |
| $K_{on,A_1}$ | $8.58 \times 0.001\ \mu m/s$ | (Goehring et al., 2011) |
| $K_{A_1A_2}$ | $10\ \mu m^4$ | This study |
| $\gamma$ | 2.0 | This study |
| $A_1^{total}$ | $1.56\ \mu m^{-3}$ | (Goehring et al., 2011) |
| $A_1^{total}$ (AB cell) | $1.56 \times 1.3\ \mu m^{-3}$ | This study |
| $A_1^{total}$ (P1 cell) | $0.988\ \mu m^{-3}$ | This study |
| $K_{off,A_1}$ | $5.4 \times 0.001\ s^{-1}$ | (Goehring et al., 2011) |
| $K_{A_1P_1}$ | $0.2\ \mu m^2/s$ | This study |
| $\alpha$ | 1.0 | (Goehring et al., 2011) |
| $D_{A_1}$ | $0.28\ \mu m^2/s$ | (Goehring et al., 2011) |
| **$P_1$ (CHIN-1)** | | |
| $K_{on,P_1}$ | $7.0 \times 0.01\ \mu m/s$ | This study |
| $P_1^{total}$ | $1.00\ \mu m^{-3}$ | This study |
| $P_1^{total}$ (AB cell) | $1.00 \times 0.3\ \mu m^{-3}$ | This study |
| $P_1^{total}$ (P1 cell) | $1.8556\ \mu m^{-3}$ | This study |
| $K_{off,P_1}$ | $1.4 \times 0.001\ s^{-1}$ | This study |
| $K_{P_1A_1}$ | $1.6225\ \mu m^4/s$ | This study |
| $\beta$ | 2.0 | This study |
| $D_{P_1}$ | $0.15\ \mu m^2/s$ | This study |
| **$A_2$ (PAR-3–PAR-6–PKC-3)** | | |
| $K_{on,A_2}$ | $11.5 \times 0.001\ \mu m/s$ | This study |
| $A_2^{total}$ | $10\ \mu m^{-3}$ | This study |
| $A_2^{total}$ (AB cell) | $10 \times 1.3\ \mu m^{-3}$ | This study |
| $A_2^{total}$ (P1 cell) | $6.3333\ \mu m^{-3}$ | This study |
| $K_{off,A_2}$ | $3.0 \times 0.001\ s^{-1}$ | This study |
| $K_{A_2P_2}$ | $0.0097\ \mu m^2/s$ | This study |
| $\delta$ | 1.0 | This study |
| $D_{A_2}$ | $0.28\ \mu m^2/s$ | This study |
| **$P_2$ (PAR-1–PAR-2)** | | |
| $K_{on,P_2}$ | $4.74 \times 0.01\ \mu m/s$ | (Goehring et al., 2011) |
| $P_2^{total}$ | $1.00\ \mu m^{-3}$ | (Goehring et al., 2011) |
| $P_2^{total}$ (AB cell) | $1.00 \times 0.3\ \mu m^{-3}$ | This study |
| $P_2^{total}$(P1 cell) | $1.8556\ \mu m^{-3}$ | This study |
| $K_{off,P_2}$ | $7.3 \times 0.001\ s^{-1}$ | (Goehring et al., 2011) |
| $K_{P_2A_1}$ | $2.55 \times 0.0001\ \mu m^4/s$ | This study |
| $\zeta$ | 2.0 | (Goehring et al., 2011) |
| $D_{P_2}$ | $0.15\ \mu m^2/s$ | (Goehring et al., 2011) |
| **Others** | | |
| $\psi$ | $0.174\ \mu m^{-1}$ | (Goehring et al., 2011) |
| $L$ | $134.5\ \mu m$ | (Goehring et al., 2011) |
| $L$ (AB cell) | $134.5 \times 0.55\ \mu m$ | This study |
| $L$ (P1 cell) | $134.5 \times 0.45\ \mu m$ | This study |

Table S5. Parameter values used in the simulations of cortical PAR polarity in $P_2$ overexpression condition

| Parameter | Value | Source |
|---|---|---|
| $A_1$ (CDC-42–PAR-6–PKC-3) | | |
| $\boldsymbol{A_1^{total}}$ **(AB cell)** | $1.56 \times 1.8\ \mu\ m^{-3}$ | This study |
| $\boldsymbol{A_1^{total}}$ **(P1 cell)** | $0.0347\ \mu\ m^{-3}$ | This study |
| $P_1$ (CHIN-1) | | |
| $\boldsymbol{P_1^{total}}$ **(AB cell)** | $1.0\ \mu\ m^{-3}$ | This study |
| $\boldsymbol{P_1^{total}}$ **(P1 cell)** | $1.0\ \mu\ m^{-3}$ | This study |
| $A_2$ (PAR-3–PAR-6–PKC-3) | | |
| $\boldsymbol{A_2^{total}}$ **(AB cell)** | $10\ \mu\ m^{-3}$ | This study |
| $\boldsymbol{A_2^{total}}$ **(P1 cell)** | $10\ \mu\ m^{-3}$ | This study |
| $P_2$ (PAR-1–PAR-2) | | |
| $\boldsymbol{P_2^{total}}$ | $16.5\ \mu\ m^{-3}$ | This study |
| $\boldsymbol{P_2^{total}}$ **(AB cell)** | $16.5\ \mu\ m^{-3}$ | This study |
| $\boldsymbol{P_2^{total}}$**(P1 cell)** | $16.5\ \mu\ m^{-3}$ | This study |

Table S6. Parameter values used in the simulations of cortical PAR polarity for zygote and daughter cells in *cye-1(RNAi)* condition

| Parameter | Value | Source |
|---|---|---|
| | $A_1$ (CDC-42–PAR-6–PKC-3) | |
| $A_1^{total}$ (AB cell) | $1.56\ \mu\mathrm{m}^{-3}$ | This study |
| $A_1^{total}$ (P1 cell) | $1.56\ \mu\mathrm{m}^{-3}$ | This study |
| | $P_1$ (CHIN-1) | |
| $P_1^{total}$ (AB cell) | $1.0\ \mu\mathrm{m}^{-3}$ | This study |
| $P_1^{total}$ (P1 cell) | $1.0\ \mu\mathrm{m}^{-3}$ | This study |
| | $A_2$ (PAR-3–PAR-6–PKC-3) | |
| $A_2^{total}$ (AB cell) | $10\ \mu\mathrm{m}^{-3}$ | This study |
| $A_2^{total}$ (P1 cell) | $10\ \mu\mathrm{m}^{-3}$ | This study |
| | $P_2$ (PAR-1–PAR-2) | |
| $P_2^{total}$ (AB cell) | $1.5\ \mu\mathrm{m}^{-3}$ or $0.3\ \mu\mathrm{m}^{-3}$ | This study |
| $P_2^{total}$ (P1 cell) | $1.5\ \mu\mathrm{m}^{-3}$ or $0.3\ \mu\mathrm{m}^{-3}$ | This study |
| | others | |
| $L$ (AB cell) | $134.5 \times 0.5\ \mu\mathrm{m}$ | This study |
| $L$ (P1 cell) | $134.5 \times 0.5\ \mu\mathrm{m}$ | This study |
